# Floral Stage Optimization and Immune Evasion Enhance *Agrobacterium*-Mediated Genome Editing in *Arabidopsis*

**DOI:** 10.1101/2025.05.07.652770

**Authors:** Mao-Sen Liu, Teng-Kuei Huang, Yi-Chieh Wang, Si-Chong Wang, Chih-Hang Wu, Chih-Horng Kuo, Erh-Min Lai

## Abstract

- *Agrobacterium*-mediated transformation via floral inoculation (AMT-FI) enables genetic engineering without tissue culture. It is widely used in the model plant *Arabidopsis thaliana*, yet its efficiency and broader applicability remain limited.
- Here, we used a dual-reporter system (RUBY and hygromycin resistance) to identify key floral stages and engineered *Agrobacterium* strains to evade plant immunity, leading to enhanced transient expression and genome editing.
- We determined that flowers opened at 6 day-post-inoculation (DPI) are optimal for high transformation efficiency, with nearly 100% of siliques harboring transformants. However, *Agrobacterium* infection induced ovule abortion, particularly in wild-type (Col-0) plants, whereas *efr* mutants lacking the EF-Tu receptor (EFR)-mediated pattern-triggered immunity (PTI) showed reduced ovule abortion. Notably, *efr* mutants exhibited more RUBY-positive ovules and significantly enhanced genome editing efficiency. Two engineered stealth *Agrobacterium* strains (AS201 and AS202) expressing a chimeric EF-Tu for evading recognition by EFR enhanced both transient transformation and genome editing efficiency. Remarkably, genome-edited T1 plants could be recovered based on phenotype or direct sequencing without the need for antibiotic selection when targeting flowers opened at 6 DPI.
- By integrating floral stage selection, immune evasion, and *Agrobacterium* engineering, this study provides a practical and versatile platform to advance plant genome engineering.

## INTRODUCTION

*Agrobacterium* species are unique bacteria capable of cross-kingdom gene transfer, leading to diseases like crown gall or hairy root in a wide range of plant species (Hwang *et al*., 2017; Lacroix & Citovsky, 2019). This remarkable ability has been harnessed in plant genetic engineering by replacing the oncogenes within T-DNA region of tumor-inducing plasmids with genes of interest (Suzuki *et al*., 2009; Hooykaas, 2023). *Agrobacterium-*mediated transformation (AMT) has become a cornerstone technique in plant biology and the development of genetically modified organisms (GMOs) that stably express agricultural traits for crop improvements (Bevan, 1984; Gelvin, 2003).

Beyond enabling stable gene integration, AMT also supports transient expression of genome-editing machinery, allowing precise genome modifications. Since the T-DNA containing these editing components can be eliminated via chromosome segregation in subsequent generations, this strategy mitigates food safety concerns associated with GMOs (Pixley *et al*., 2022).

*Agrobacterium* naturally infects somatic plant cells by responding to wound signals in roots or stems. Interestingly, it can also target germline tissues such as female gametophytes, which laid the foundation for the floral dip method widely used in *Arabidopsis thaliana* (Clough & Bent, 1998; Angulo *et al*., 2023). This method, referred to here as *Agrobacterium-*mediated transformation by floral inoculation (AMT-FI), bypasses labor-intensive tissue culture, offering a more efficient transformation pathway. However, its application in species beyond *A. thaliana* has limited success (Bélanger *et al*., 2024).

Although many plant factors that promote or suppress tumorigenesis and AMT have been identified, the genes involved in AMT-FI may be distinct (Hwang *et al*., 2017; Lacroix & Citovsky, 2019; Weisberg *et al*., 2023). Several *Arabidopsis* mutants or ecotypes resistant to root transformation remain susceptible to AMT-FI (Nam *et al*., 1997; Mysore *et al*., 2000). Notably, *polQ* mutants lacking DNA polymerase theta are deficient in T-DNA integration via AMT-FI, although their capability for stable transformation in roots remains debated (van Kregten *et al*., 2016; Nishizawa-Yokoi *et al*., 2021). These findings highlight that developmental processes may impose different requirements for successful transformation in germline and somatic cells. Understanding the genetic and molecular basis of AMT-FI is crucial to expanding its utility across plant species.

Plant immunity plays a pivotal role in suppressing tumorigenesis and AMT efficiencies (Tiwari *et al*., 2022; Dodds *et al*., 2024). Upon *Agrobacterium* infection, plants recognize pathogen-associated molecular patterns (PAMPs) such as EF-Tu, flagellin, and cold-shock proteins by pattern-recognition receptors (PRRs), activating pattern-triggered immunity (PTI) that suppress transformation (Zipfel *et al*., 2006; Wu *et al*., 2014; Saur *et al*., 2016; Wang *et al*., 2018; Fürst *et al*., 2020). Salicylic acid (SA) is a key defense hormone that reduces tumor formation and transient transformation (Anand *et al*., 2008). Additionally, antimicrobial secondary metabolites, including glucosinolates and camalexin, restrict *Agrobacterium* infection at early and late stages, respectively (Shih *et al*., 2018). These defense strategies collectively highlight the resilience of plants to counteract *Agrobacterium* infection and limit its transformation success.

In *Arabidopsis*, the EF-Tu receptor (EFR) plays a key role in recognizing *Agrobacterium* EF-Tu and inhibiting transient transformation (Zipfel *et al*., 2006; Wu *et al*., 2014; Wang *et al*., 2018). However, EFR does not exhibit a significant impact on stable transformation or gene targeting efficiency (van Tol *et al*., 2022; Yang *et al*., 2023). To date, its influence on AMT-FI-mediated genome editing has not been fully elucidated.

In this study, we investigated the temporal and spatial localization of *Agrobacterium* cells and transformation events during AMT-FI in *A. thaliana.* A dual reporter system, combining a visible RUBY marker with a hygromycin resistance marker, was utilized to identify the floral tissues and stages most susceptible to AMT. Furthermore, we examined the impact of EFR-mediated PTI in transformation and genome-editing efficiencies. Our results revealed that AMT events peak in flowers opened at 6 day-post-inoculation (DPI), with *efr* mutants exhibiting significantly higher genome editing efficiency than Col-0. Moreover, we engineered two independent *Agrobacterium* strains expressing a chimeric EF-Tu, which further enhanced both transient expression and genome editing efficiency without significantly altering stable transformation frequency. These findings offer new strategies to facilitate AMT-FI-based genome editing in plants.

## MATERIALS AND METHODS

### Plant materials and growth condition

*Arabidopsis thaliana* Col-0, EF-Tu receptor (EFR) mutants, *efr*-1 (SALK_044334) and *efr*-2 (SALK_068675) were used in this study. Seeds were sown in soil (peatmoss:vermiculite:perlite = 12:1:1), germinated, and grown in a growth room under conditions of 22 °C with 16h light/8h dark photoperiod.

### Bacterial growth condition

For tracking *Agrobacterium* localization and transformation events, *Agrobacterium* cells were cultured in 523 with appropriate antibiotics (Kado & Heskett, 1970). For genome editing experiments, LB medium supplemented with appropriate antibiotics was used. Freshly streaked out *Agrobacterium* colonies grown on 523 or LB agar were cultured in the same medium (5 mL) for 2 days at 28 °C. The cultures were subsequently diluted 1:1000 in 50 mL 523 or LB, grown for 22-24 hours, harvested by centrifugation at 6000x g for 20 minutes. The cell pellets were resuspended in infiltration medium (1/2 MS, 5% sucrose, 0.04% silwet-L77, 44 nM BAP and 200 μM acetosyringone, pH=5.7) to OD_600_=1.

### Plasmid construction

Plasmid and primer information are listed in Table S1 and S2, respective. 35S::RUBY plasmid (Addgene#160908) which provided a visible RUBY reporter and hygromycin resistance (Hyg^R^) reporter was obtained from Addgene.

The gentamycin resistance gene (Gen^R^) of pRL662-GFP(S65T) (Yu *et al*., 2021) was replaced by spectinomycin resistance gene from 35S::RUBY to obtain pRL662- GFP(S65T)/Spe^R^ by use of Gibson Assembly® Master Mix (New England Biolabs, USA). The expression of mCherry2 with a 3’-end nuclear localization signal (mCherry-NLS) was generated from pEB2-2ndVal mCherry2-L (Addgene#104027) by PCR. PCR was performed by use of KAPA HiFi HotStart ReadyMix PCR Kit (KapaBiosystems, South Africa) and PCR products were cloned into the pJet1.2/blunt plasmid following the user manual of CloneJET PCR Cloning Kit (ThermoFisher Scientific, USA). 35S::mCherry2-SV40/Kan^R^ binary vector was constructed with Golden Gate Assembly.

A3A-CBE coding region was amplified from pYPQ265E2 (Addgene #164719) and was subcloned into XbaI/SacI digested pHEE901 (Addgene #91707) via Gibson Assembly. Protospacers were cloned into pA3A-CBE vectors as previously described (Steinert *et al*., 2015). To generate the *tufAB^Pst4-15^* mutants in GV3101, we employed double crossover vector using pEML041003-tufAPst4-15 and pEML41003- tufBPst4-15. Two 2.5-kb DNA fragments of modified *tufA* and *tufB* were amplified by two pairs of primers, and cloned into pEML041003 by BsaI digestion. AS201 was generated by modifying *tufB* followed by *tufA*, while AS202 was *tufA* followed by *tufB*. In both constructs, the amino acids at positions 4-15 of EF-Tu were substituted from the native sequence SKFERNKPHVNI to the *Pseudomonas syringae* variant EKFDRSLPHCNV.

### Seedling transient transformation and GUS activity assays

*A. thaliana* seedling transient expression assay was performed by the AGROBEST (*Agrobacterium*-mediated enhanced seedling transformation) method as previously described, with minor modification (Wu *et al*., 2014). *Agrobacterium* cultures were adjusted to an OD_600_=0.01 in AB-MES medium for co-culture with seedlings. Infected seedlings were collected for GUS activity assays after 2.5 day-post-inoculation.

### *Agrobacterium*-mediated transformation by floral inoculation (AMT-FI)

AMT-FI was performed according to the published procedure (Clough & Bent, 1998) with modifications. For tracking *Agrobacterium* cells and transformation events, each *Arabidopsis* plant was grown in a small pot (5 cm upper diameter x 5 cm high x 3.5 cm bottom diameter). For genome editing experiments, three plants were grown in a large pot (7.8 cm upper diameter x 7.8 cm high x 5.7 cm bottom diameter). Plants were grown until they reached ∼5 weeks old, with primary inflorescences reaching 10 to 15 cm tall. The primary inflorescence was then removed to promote the emergence of secondary inflorescences over the next 5 to 6 days. Immediately prior to floral dipping, all open flowers were removed. Inflorescences were submerged in *Agrobacterium*-containing infiltration medium for 40 seconds, then plants were laid horizontally in trays, covered with plastic film to maintain high humidity, kept in the dark and incubated for 20 to 24 hrs. Plants were returned to the growth room under the same growth condition.

For time-course experiments, newly opened flowers were labeled daily following inoculation. GFP-labeled *Agrobacterium* cells and mCherry2 T-DNA reporter signals were examined in ovaries using a Zeiss LSM 510 Meta confocal microscope RUBY signals were monitored visually in plants transformed with GV3101 carrying the 35S::RUBY construct, assisted by a stereo microscope. A Zeiss Lumar V12 stereo fluorescent microscope was used to capture images of floral organs, developing ovules, seeds, and seedlings.

To evaluate AMT-FI efficiency by RUBY marker, siliques from RUBY-transformed plants were collected daily. To assess stable transformation efficiency, seeds were sterilized in 70% ethanol for 10 min and plated on a 1/2 MS medium supplemented with 25 mg/L hygromycin. The number of germinated seedlings and Hyg^R^ seedlings was recorded. AMT-FI efficiency was determined based on the proportion of Hyg^R^ seedlings relative to the total number of germinated seedlings.

### Assessment of genome editing efficiency

To simultaneously measure the genome editing and AMT-FI efficiency, seeds from plants transformed with *Agrobacterium* carrying a binary vector with T-DNA-encoded base editor, a sgRNA and a hygromycin resistance gene were surface sterilized and germinated on a 1/2 MS medium supplemented with either 5 mg/L tribenuron or 25 mg/L hygromycin, both containing 100 mg/L timentin. After seven days, the numbers of tribenuron-resistant, hygromycin-resistant, and albino T1 seedlings were recorded. AMT-FI efficiency was estimated using a weight-based approach, with 10 mg of seeds estimated to contain approximately 500 seeds.

### Statistical analysis

An unpaired two-tailed Student’s *t-*test, performed using GraphPad Prism 10, was used to assess statistical differences either between Col-0 WT and *efr* mutants or across time-course datasets.

## RESULTS

### *Agrobacterium* enters the female reproductive organ within one hour but transformation events occur in open flowers after five day-post inoculation

Although transformation is thought to occur when the female gametophyte becomes accessible, direct links between floral stage and transformation success remain unclear. To investigate the spatial and temporal dynamics of *Agrobacterium* infection and transformation, we engineered a fluorescently labeled *Agrobacterium* strain for live tracking. *Agrobacterium* GV3101 carrying the disarmed Ti plasmid pMP90 (hereafter referred as GV3101), constitutively expressing GFP marker and a T-DNA expressed mCherry2, was used for floral inoculation in *A. thaliana* Col-0 (Fig.1a, 1b). GFP-expressing *Agrobacterium* cells were observed as early as 1 hour-post-inoculation (HPI) in the early developmental stage of open gynoecia, the female reproductive organ lacking central placenta tissue (Fig. 1c) but were not detected in the later stage of gynoecia containing central placenta tissue (Fig. 1d). This implied that developing placenta may act as a physical barrier to *Agrobacterium* entry during early infection. By 6 HPI, *Agrobacterium* cells accumulated in the locule of open gynoecia (Fig. 1e, 1f), whereas no signal was seen in closed gynoecia (Fig. 1g). This indicated that open gynoecia serve as the main entry point of *Agrobacterium* into the locule before development of female gametophyte.

**Figure 1.**
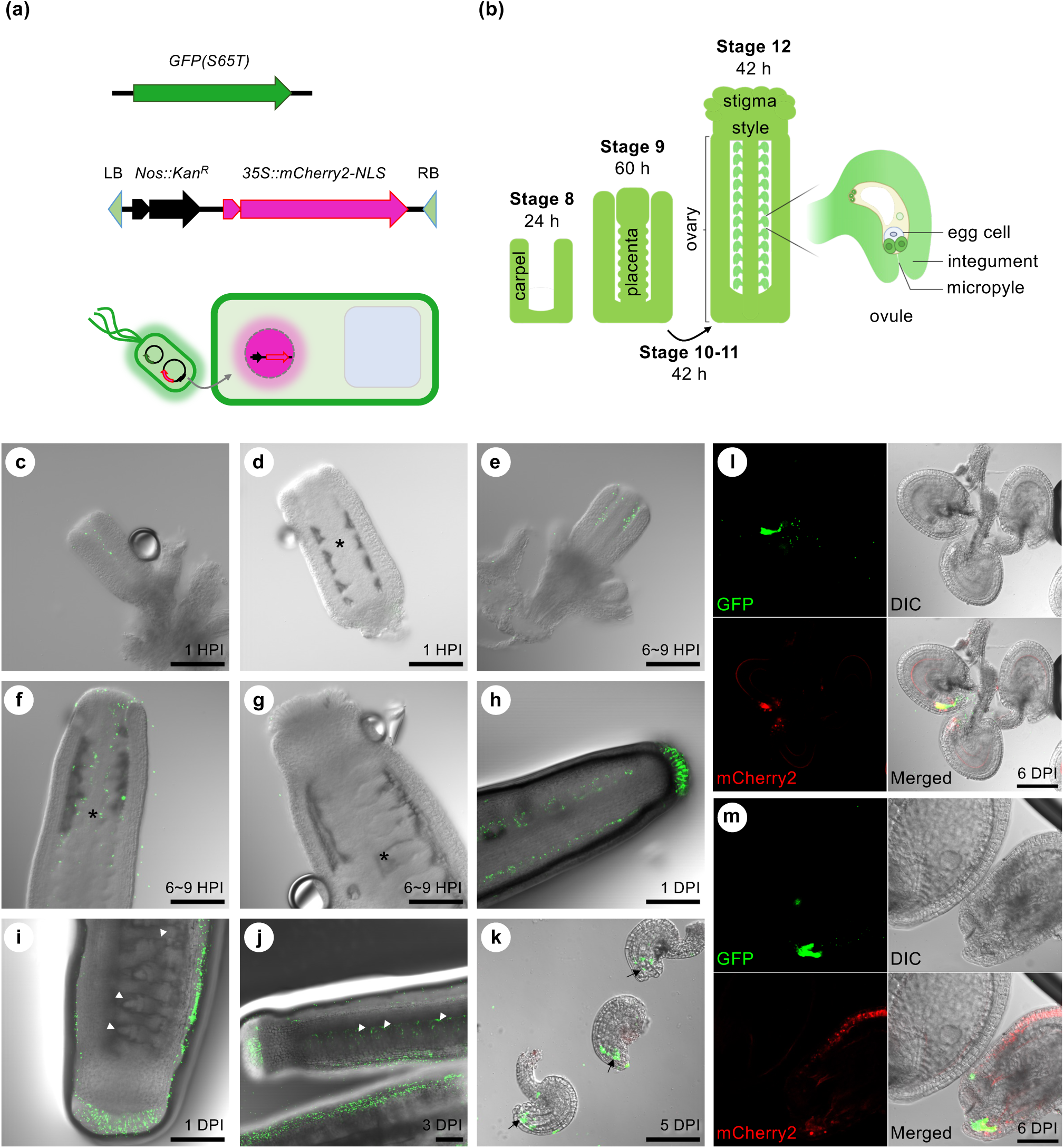
*Agrobacterium* enter open gynoecium within 1 hour but transformation events occur in open flowers after 5 day-post inoculation. (a) *Agrobacterium* GV3101 carrying pRL662-GFP(S65T)/Spe^R^ and the binary vector 35S::mCherry2-NLS was used for floral inoculation on *Arabidopsis thaliana* Col-0 to visualize *Agrobacterium* localization and transformation events. (b) Schematic illustration of the structure and development of gynoecium from early to mature stage with stigma, style, ovary containing ovules. The floral stages and their corresponding durations were indicated according to Smyth et al. The diagram was created with BioRender.com. After one hour-post-inoculation (HPI), *Agrobacterium* cells had already entered the locule of developing gynoecium (c) but were absent in later-stage gynoecia containing developed placenta tissue (d). Between 6-9 HPI, more *Agrobacterium* cells were observed in the locule (e), even at it began to close (f). *Agrobacterium* was not detected in fully closed gynoecia (g). At one day-post-inoculation (DPI), *Agrobacterium* cells were detected in young closed ovaries (h) but not in older ovaries undergoing ovule development (i). At 3 DPI (j), *Agrobacterium* cells persisted in ovaries at late developmental stages. By 5 DPI, *Agrobacterium* cells were enriched around the micropyle (arrow) (k), though no mCherry2 signal was detected. At floral open stage after 6 DPI, *Agrobacterium* accumulated around the micropyle, and mCherry2 signal became visible (l and m). Asterisks (*) indicate placenta tissue; arrowheads denote developing ovule; arrows indicate *Agrobacterium* located around micropyle. Panels (a to k) were merged channel images; panels (I and m) show GFP, mCherry2, DIC and merged views. Scale bar= 100 µm.

At one day-post-inoculation (DPI), *Agrobacterium* cells were detected in locules of younger gynoecia (Figs. 1h, S1a-f) but were absent in older gynoecia undergoing ovule development (Figs. 1i, S1g-n). Among more than 30 gynoecia examined, all 13 open gynoecia observed at 6 HPI and 1 DPI showed detectable GFP signals in the locule, indicating the entrance of *Agrobacterium* prior to the closure of gynoecia. At 3 DPI, *Agrobacterium* persisted in the locules of more mature gynoecia (Fig. 1j), and at 5 DPI, just prior to floral opening (anthesis), *Agrobacterium* cells were enriched around the micropyles, the ovular opening through which the pollen tube and possibly *Agrobacterium* gain access (Fig. 1k). Despite the sustained presence of *Agrobacterium*, successful T-DNA expression indicated by mCherry2 fluorescence was not detected before anthesis (Fig. 1c-k). Similarly, the RUBY reporter signal was absent in gynoecial tissues until after flower opening (Fig. 2c). Following anthesis, mCherry2 signals became visible in cells adjacent to the egg cell/zygote and in the micropyle region where *Agrobacterium* cells had accumulated (Fig. 1l, 1m). Transformation signals were also detected in the integument, the ovule’s outer protective layer (Fig. 1m).

**Figure 2.**
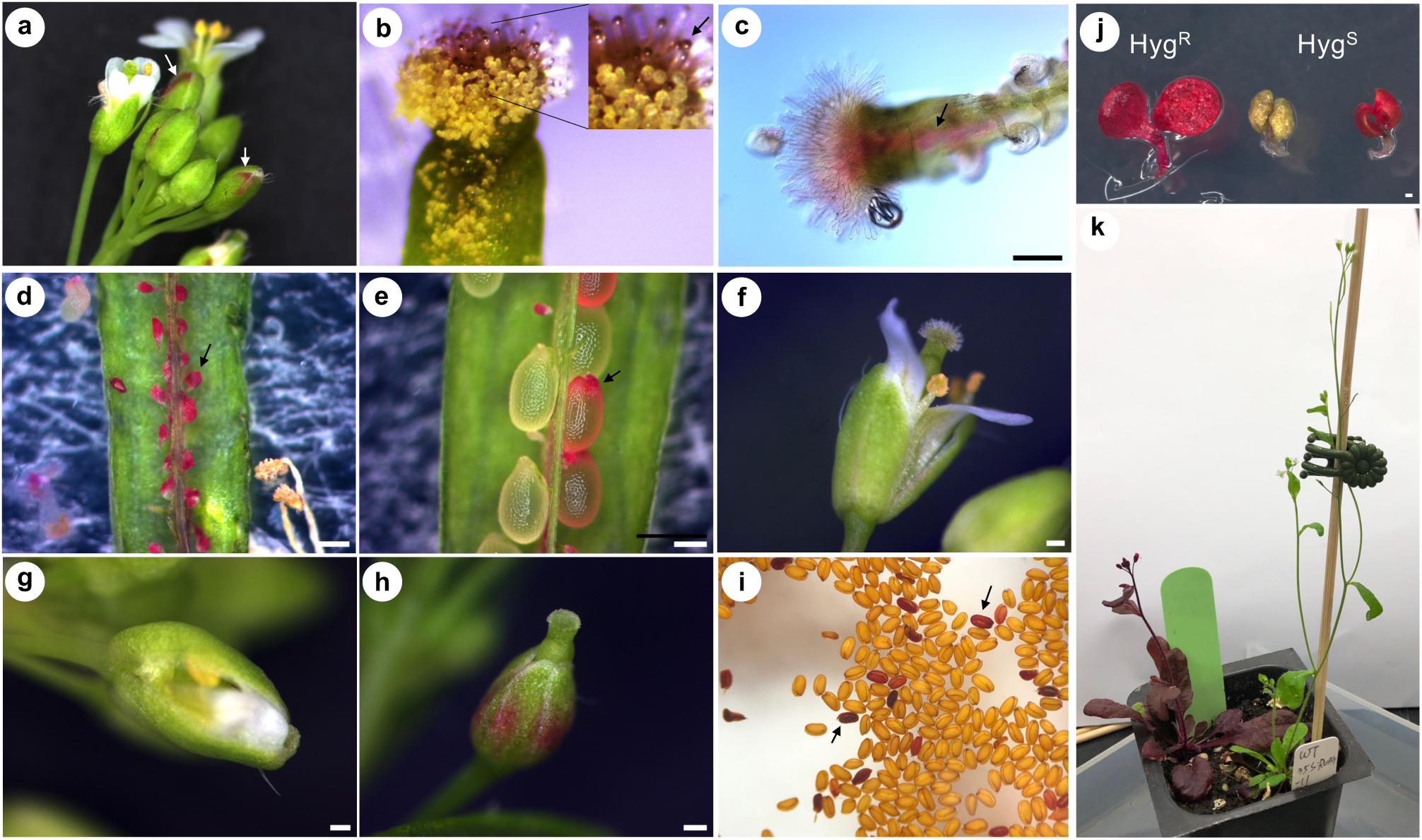
RUBY marker revealed transformation events in floral organs and female tissues. RUBY signals were observed in flowers (a), specifically on the stigma but not on pollen grains (b), and in the transmitting tissue (c). In siliques, RUBY expression was detected in both abortive (d) and developing ovules (e). Flower morphology after AMT-FI ranged from normal open flowers (f) to partially open flowers (g), and flowers with only gynoecia emerging from sepals (h). RUBY signal intensity varied among seeds (i) and plants (j, k). (j) At the seedling stage, hygromycin-resistant (Hyg^R^) and RUBY-positive seedling (left), non-transgenic seedling (middle), and RUBY-positive but hygromycin sensitive (Hyg^S^) seedling (right) were observed. (k) Transgenic T1 plants with variable RUBY expression patterns from strong whole-plant expression to undetectable signal. Scale bar = 200 µm.

Together, these observations reveal that while *Agrobacterium* rapidly enters the open gynoecium as early as 1 HPI, transformation events are temporally restricted, becoming detectable only in flowers that opened from 5 DPI onward. This correlates with key developmental transitions in ovule accessibility and gametophyte maturity.

### RUBY-based reporter system uncovers transformation dynamics and floral abnormalities

The current method for quantifying transformation efficiency of AMT-FI primarily relies on the use of antibiotic resistance genes, using survival rates of T1 seedlings as a proxy for transformation efficiency. However, this approach overlooks transient transformation events that occur at early stages post-inoculation. To better capture both the transient and stable transformation events, we developed a dual-reporter system combining the visible RUBY marker with a hygromycin-resistance gene (Fig. 3a).

**Figure 3.**
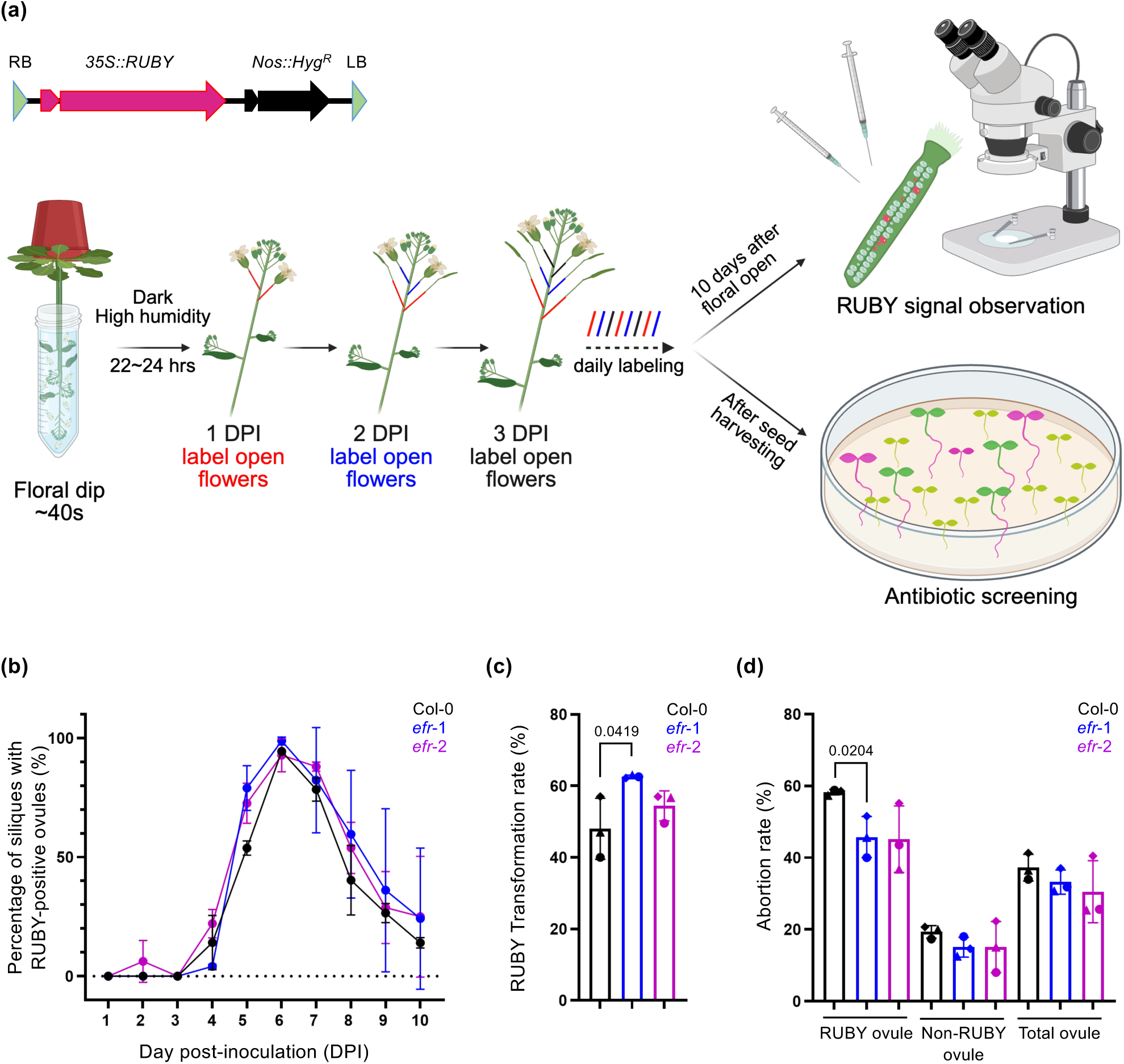
*Agrobacterium* infection and EFR-mediated immunity exert negative impacts on ovule development. (a) Experimental design: a schematic diagram illustrates the 35S::RUBY construct and experimental workflow. Transformation efficiency was evaluated based on the presence of RUBY-positive ovules and hygromycin-resistant seedlings. The diagram was created with BioRender.com. (b) Percentage of siliques with RUBY-positive ovules recorded daily for Col-0 WT, *efr*-1, and *efr*-2 from flowers that opened at 1-10 DPI. Data are mean ± SD of three biological replicates from two independent experiments. (c) RUBY-based transformation. Percentage of RUBY-positive ovules (include both developed and abortive ovules) per silique collected from flowers of Col-0 WT, *efr*-1 and *efr*-2 opened at 5-9 DPI was calculated (all data points shown in Fig. S3a). Data are mean ± SD of average percentage of RUBY-positive ovules from three independent experiments (circle, triangle and rhombus symbols). Statistically significant differences (*p* < 0.05) between Col-0 WT and *efr* mutants are indicated. (d) Ovule abortion rate. The abortion rate of RUBY-positive, non-RUBY, and total ovules per silique collected from flowers of Col-0 WT, *efr*-1 and *efr*-2 opened at 5-9 DPI was calculated (all data points shown in Fig. S3c). Data are mean ± SD of average abortion rate from three independent experiments (circle, triangle and rhombus symbols). Statistically significant differences (*p* < 0.05) between Col-0 WT and *efr* mutants are indicated.

We first assessed the floral organs exhibiting RUBY expression. The earliest detection of RUBY signals was observed at 4 DPI, localized at the sepals of floral buds (Fig. 2a). By 5 DPI, RUBY expression appeared on the stigma of open flowers but not on pollen grains (Fig. 2b). Dissection of the ovaries revealed RUBY signal in the transmitting tissues (Fig. 2c), as well as in ovules and developing seeds (Fig. 2d, 2e). While many RUBY ovules were abortive (Fig. 2d), others continued to develop normally (Fig. 2e). Most flowers developed and opened normally (Fig. 2f) while a minor subset exhibited developmental abnormalities. These included flowers with partially open petals (Fig. 2g) and those where only the gynoecium emerged from the sepals (Fig. 2h). In mature seeds, RUBY expression ranged in intensity from light pink to dark purple (Fig. 2i), indicating variable expression levels.

In the hygromycin resistance assay, most Hyg^R^ seedlings only showed weak or sparse RUBY signals, with ∼10% or less of T1 seedling showing RUBY in whole seedlings (Fig. S2). Because hygromycin-sensitive seedlings can continue to grow for up to seven days before growth arrest due to their inability to form roots on hygromycin-containing medium, we can still observe RUBY⁺/Hyg^S^ seedlings, although their frequency is low (less than 1% of total transformants, Fig. 2j).

Consequently, not all hygromycin-resistant plants exhibited uniform RUBY expression throughout the plants (Fig. 2k). Overall, the percentage of RUBY-positive seeds was higher than the frequency of Hyg^R^ seedlings. This discrepancy may reflect transient expression or expression of RUBY marker in the integument, which does not necessarily indicate stable T1 transformation.

Collectively, the dual-reporter system offers a more comprehensive and sensitive tool for monitoring transformation events across the reproductive tissues. The RUBY reporter revealed that AMT-FI enables transformation in a variety of tissues within the female reproductive organ. However, it also uncovered associated developmental abnormalities in flowers and ovules, which in some cases prevent transformed ovules from developing into viable seeds.

### *Agrobacterium* infection and EFR-mediated immunity synergistically disrupt ovule development and seed viability

Since transformation events were only observed in ovaries after anthesis, we sought to trace the occurrence of AMT events in developing siliques using RUBY reporter as a marker (Fig. 3a). Flowers that opened between 1 to 10 DPI were labeled daily.

Once embryos were fully developed, approximately 10 days after floral opening and pollination (Baud *et al*., 2002), each corresponding silique was examined for RUBY signals in ovules/developing seeds.

To assess the impact of EFR-mediated immunity on AMT-FI efficiency, we compared transformation efficiency in Col-0 wild-type (WT) plants and two *EFR* loss-of-function mutants (*efr*-1 and *efr*-2) (Zipfel et al., 2006). Transformation events were primarily detected in flowers that opened between 5 to 9 DPI, with the highest frequency, nearly 100% of siliques showing RUBY-positive ovules in flowers opened at 6 DPI in both Col-0 WT and *efr* mutants (Fig. 3b). The percentage of siliques harboring RUBY ovules did not differ significantly between WT and *efr*, suggesting that *Agrobacterium* can access the open gynoecium and subsequently transform ovules with comparable success across genotypes. However, the proportion of RUBY-positive ovules per silique was moderately higher in *efr* mutants, suggesting that compromising EFR-mediated PTI can enhance transformation efficiencies (Figs. 3c, S3a, S3b). Importantly, the number of siliques per plant and total seed number did not show significant difference between WT and *efr* mutants (Fig. S4). Thus, the higher RUBY transformation rate per silique in *efr* mutants (Figs. 3c, S3b) is not caused by variation of siliques or seeds between Col-0 and *efr* mutant plants.

Given that AMT-FI often induced floral abnormalities and ovule abortion, we further investigated whether EFR-mediated PTI affects ovule viability. Strikingly, in Col-0 WT, nearly 60% of RUBY-positive ovules were abortive, indicating a strong negative impact of AMT-FI on ovule development. In *efr* mutants, although ovule abortion still occurred, the rate was significantly reduced to ∼45%. In contrast, fewer than 20% of non-RUBY ovules were abortive, with no significant difference between Col-0 WT and *efr* mutants (Fig. 3d). On average, *Agrobacterium* infection resulted in an ovule abortive rate of ∼35% (Figs. 3d, S3c).

Notably, a strong negative correlation was observed between transformation rate and seed viability. Flowers that opened at 6 DPI, despite showing the highest AMT- FI efficiency, produced low number of viable seeds (Fig. 4a), likely due to the high abortion rate of RUBY-positive ovules (Fig. 3d). However, *efr* mutants produced significantly more viable seeds per silique than Col-0 WT from flowers that opened at 6 to 8 DPI (Fig. 4a), supporting a role for EFR-mediated immunity in transformation-associated ovule abortion.

**Figure 4.**
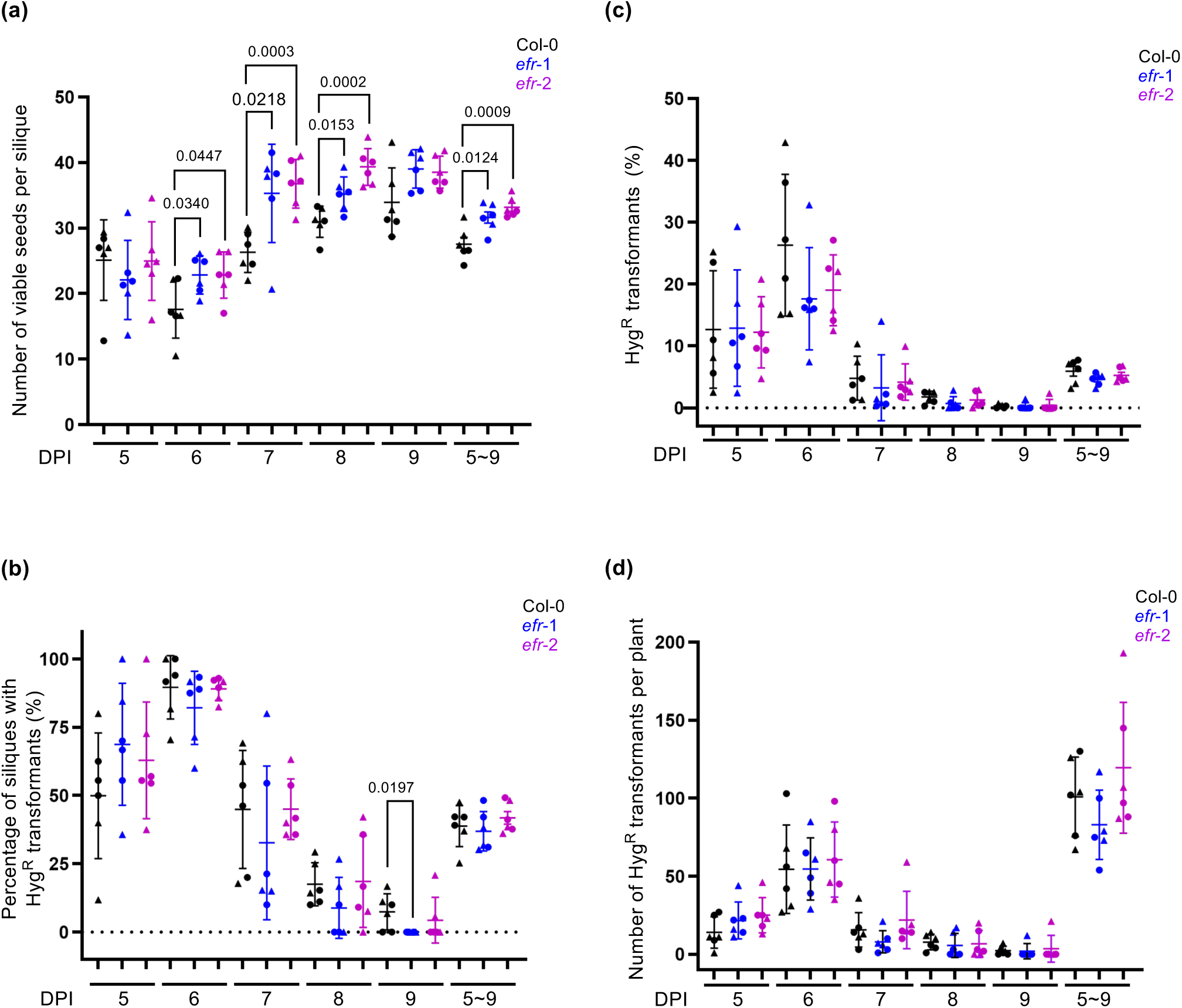
AMT events occur mainly in flowers opened at 5-9 DPI, peaking at 6 DPI. Siliques and seeds developed from flowers that opened at 5-9 DPI in Col-0 WT, *efr*-1 and *efr*-2 mutants were analyzed through viability (a) and hygromycin resistance (Hyg^R^) screening (b, c, d) and viability (d). Data are (a) average number of viable seeds per silique, (b) percentage of siliques containing Hyg^R^ transformants, (c) percentage of Hyg^R^ transformants and (d) number of Hyg^R^ transformants per plant. Data are mean ± SD of six biological replicates from two independent experiments (circle and triangle symbols). Statistically significant differences (*p* < 0.05) between Col-0 WT and *efr* mutants are indicated.

To determine whether the increased number of viable seeds per silique in *efr* mutants is a consequence of AMT-FI, we quantified seed production in plants treated with infiltration medium, and those inoculated with *Agrobacterium* GV3101 (Fig. S5). Under both conditions, *efr* mutants produced more seeds per silique than Col-0 WT (Fig. S5a, S5b). However, *Agrobacterium* infection further reduced seed numbers in both genotypes. To assess seed viability, we evaluated germination rates. Seeds from plants treated with infiltration medium germinated at nearly 100% but the germination rate is reduced in both Col-0 and *efr* (∼90%) with *Agrobacterium* infection. However, when the number of developed/germinated seeds following *Agrobacterium* infection is normalized to that of seeds treated with infiltration medium, *efr* mutants exhibited higher seed development and germination rates (Fig. S5a).

Taken together, the results suggested that the elevated abortion rates and reduced seed viability observed here are largely attributable to *Agrobacterium* infection because ovule abortion rates are typically less than 1% under normal growth conditions (van Tol *et al*., 2022). EFR-mediated immunity may also reduce seed development and viability but pathogen-induced stress from *Agrobacterium* infection has a more substantial negative impact than through EFR alone. While alleviation of EFR-mediated immunity reduced the abortion rate among transformed ovules by ∼15%, it did not restore ovule viability to levels observed in non-transformed ovules (Fig. 3d). These findings suggest that, in addition to EF-Tu PAMP recognition, other *Agrobacterium*-derived factors likely contribute to ovule developmental disruption during AMT-FI.

### Temporal profiling reveals the optimal window for stable AMT-FI transformation

Although the RUBY-positive ovules and seeds indicate transformation events, not all develop into viable transgenic T1 seeds. To more accurately evaluate inherited transformation efficiency, we quantified the number of viable transgenic seeds per silique and stable transformation rate using hygromycin resistance as a selection marker. Stable transformation efficiency was calculated based on the number and percentage of Hyg^R^ seedlings collected daily from flowers opened at 5-9 DPI (Figs. 4, S4c).

As expected, siliques from flowers opened at 6 DPI exhibited the highest transformation efficiency, with an average of ∼90% and a peak at 100% of siliques containing one or multiple Hyg^R^ transformants (Fig. 4b). Consistently, stable transformation efficiency peaked at this time point, with an average of ∼26% and a peak at 40% of T1 progeny exhibiting hygromycin resistance (Fig. 4c). The number of T1 Hyg^R^ transformants per plant also reached its highest at 6 DPI, averaging ∼55 and peaking at 100 per plant (Fig. 4d).

These results highlight that flowers opened at 6 DPI as the optimal floral stage for achieving the highest frequency of stable AMT-FI-mediated transformation in *Arabidopsis*.

### EFR-mediated immunity suppresses genome editing efficiency

Notably, no significant increase in stable transformation efficiency was observed in *efr* mutants compared to Col-0 WT (Fig. 4c), similar to the previous findings (van Tol *et al*., 2022; Yang et al., 2023). Previous studies have shown that *efr* mutants exhibit enhanced *Agrobacterium-*mediated transient expression (Zipfel et al., 2006; Wu, Liu et al., 2014). Since transient expression of CRISPR/Cas9 components is sufficient to induce genome editing without stable T-DNA integration, we next investigated whether EFR-mediated immunity adversely affects gene editing efficiency during AMT-FI.

We constructed a T-DNA vector carrying dual selection markers: a hygromycin resistance gene for selecting transgenic plants and a CRISPR/Cas9 system targeting the *ALS* gene, which confers dominant herbicide resistance (Chen *et al*., 2017), for identifying genome-edited plants (Fig. 5a). Given that the egg-cell is a major target of AMT-FI (Desfeux *et al*., 2000), we adopted a strategy that enables editing in monoploid germline cells at early developmental stages. Specifically, we applied an egg-cell-specific promoter previously shown to induce heritable mutations in the T1 generation (Wang *et al*., 2015). To enhance genome editing efficiency and streamline the screening process, we employed an A3A/Y130F-CBE_V01 base editor (termed as A3A-CBE in this study), which has demonstrated efficient C-to-T base editing in both rice and *Arabidopsis* under a ubiquitin promoter (Ren *et al*., 2021).

**Figure 5.**
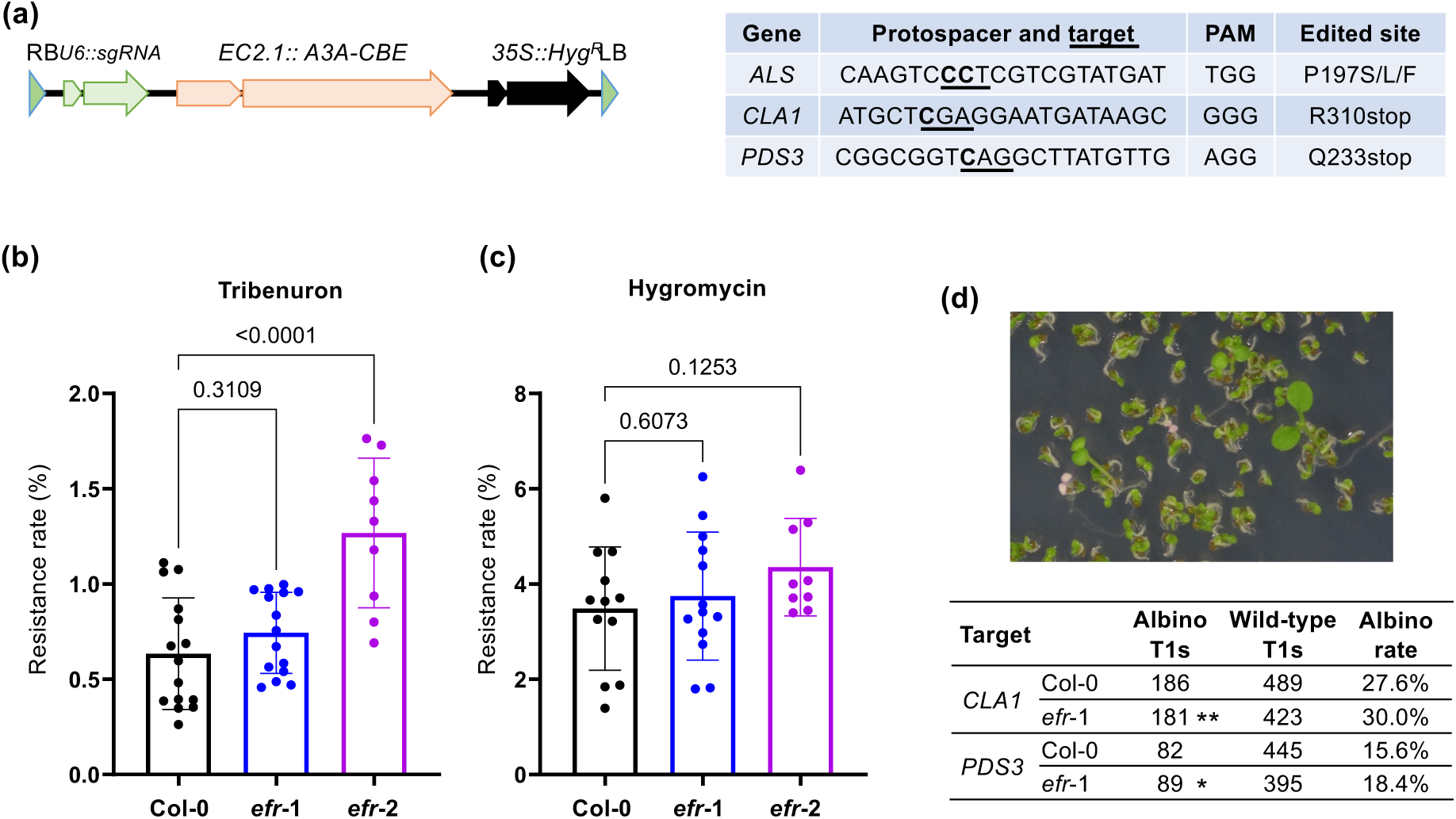
EFR suppresses genome editing efficiency. *Agrobacterium* GV3101 carrying a binary vector encoding a base editor and a sgRNA was used for floral inoculation in Col-0 WT and *efr* mutants to assess genome editing and stable transformation efficiencies. (a) A schematic diagram of the binary vector structure for base editing and target gene information. (b) Genome editing efficiency assessed by tribenuron-resistant T1 plants with targeting the *ALS* Pro197 locus. (c) Stable transformation efficiency evaluated by hygromycin resistance. (b) and (c): Seeds are collected from flowers that opened at 5-9 DPI. Each data point represents biological replicates from six independent experiments. Data are mean ± SD, with *p*-values calculated by Student’s t-test between Col-0 WT and *efr* mutants. (d) Genome editing efficiency assessed based on albino phenotype in T1 plants, resulting from by loss-of-function mutations at locus *CLA1 and PDS3.* Statistically significant differences between Col-0 WT and *efr* are indicated (\**p* < 0.05, \*\**p* < 0.01, chi-square analysis).

Using this construct, we transformed Col-0 WT and *efr* mutants and assessed T1 progeny for T-DNA integration (via hygromycin resistance) and genome editing (via tribenuron resistance conferred by *ALS* editing). The A3A-CBE system achieved an overall editing frequency of approximately 0.5% in total T1 seeds (Fig. 5b). Both *efr*-1 and *efr*-2 mutants showed higher editing rates, despite the increase being statistically significant only in *efr*-2. We also evaluated stable transformation efficiency by sowing T1 seeds on a hygromycin-containing medium but no significant increase can be observed in *efr* mutants (Fig. 5c).

To further validate these findings, we targeted *CLA1* and *PDS3*, two genes that cause albino phenotypes when biallelically edited (Liu *et al*., 2022). T1 seeds were germinated on a hygromycin-containing medium, and the numbers of green and albino seedlings were quantified (Fig. 5d). We observed a slight but consistent increase in the proportion of albino seedlings in *efr*-1 compared to Col-0 WT, supporting the conclusion that EFR-mediated immunity suppresses genome editing efficiency during AMT-FI.

### Modification of *Agrobacterium* EF-Tu enhances transient gene expression and genome editing efficiency but not stable T-DNA integration efficiency

To further validate the role of EFR in suppressing genome editing efficiency during AMT-FI, we tested whether an *Agrobacterium* strain that evades EFR recognition mediates increased genome editing efficiency. To achieve this, we engineered *Agrobacterium* to express a modified EF-Tu incapable of triggering EFR-mediated PTI. *Agrobacterium* GV3101 possesses two EF-Tu-encoding genes, *tufA and tufB*, which produce a translation elongation factor. Our initial attempts to delete both *tuf* genes were unsuccessful, likely due to their essential function. However, a recent study demonstrated that replacing the DNA fragment encoding entire or N-terminal region (amino acids 1-15) of *Agrobacterium* EF-Tu with that of *Pseudomonas syringae* pv. *tomato* (*Pst*) DC3000 reduced PTI activation and improved transient expression efficiency in *A. thaliana* leaves (Yang *et al*., 2023).

Because the first three amino acids of EF-Tu are identical between GV3101 and *Pst* DC3000, we adopted a similar concept but applied a different experimental strategy to engineer GV3101 by replacing amino acids 4-15 of both *tufA* and *tufB* with the corresponding region from *Pst* EF-Tu, referred to as *tufAB^Pst4-15^* (Fig. 6a). We generated two independent *tufAB^Pst4-15^* strains, named as AS201 and AS202, and compared them to the parental GV3101 strain for transient expression, genome editing and stable transformation efficiencies. To assess transient transformation, we transformed *A. thaliana* seedlings for transient expression analysis using a T-DNA-encoded GUS reporter (Wu *et al*., 2014). Both AS201 and AS202 showed significantly higher transient GUS activity in Col-0 WT compared to GV3101 (Fig. 6b, 6c). As expected, GV3101 exhibited ∼20-fold higher GUS activity in *efr*-1 than in Col- 0 due to the absence of EF-Tu recognition. However, in *efr*-1, AS201 and AS202 exhibited lower GUS activity than GV3101, indicating that the engineered EF-Tu proteins may compromise transformation efficiency when PTI is not a barrier for infection.

**Figure 6.**
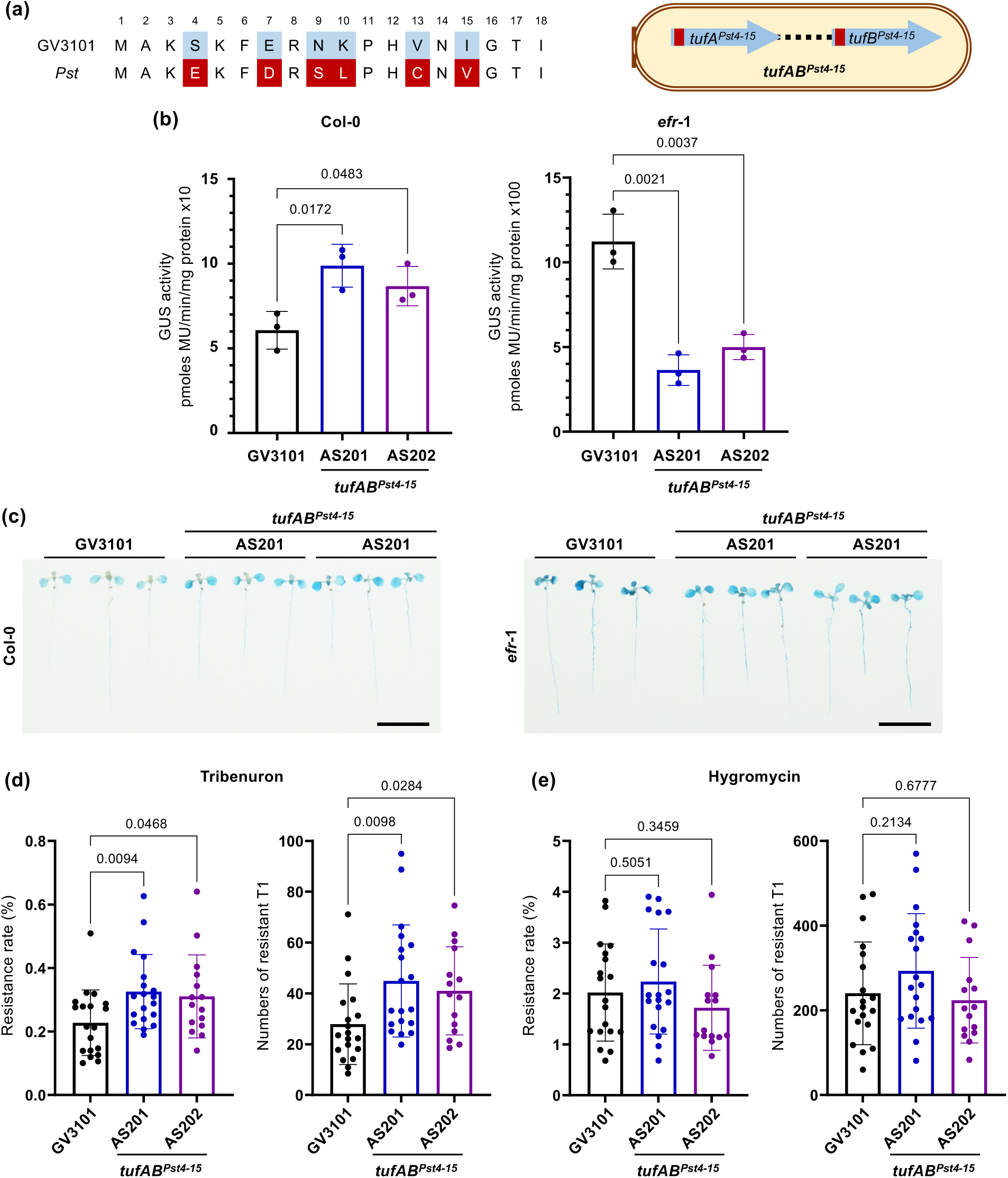
Modification of *Agrobacterium* EF-Tu enhances transient gene expression and genome editing efficiency but not stable T-DNA integration efficiency. The two engineered *Agrobacterium tufAB^Pst4-15^* strains, AS201 and AS202, were assessed for its efficiency in transient expression, genome editing, and stable transformation in comparison to the parental *Agrobacterium* GV3101. (a) A schematic diagram illustrating the difference of *tufAB* sequences at amino acids 4^th^ to 15^th^ between *Agrobacterium* GV3101 and *Pseudomonas syringae* (*Pst*). (b, c) Comparison of transient transformation efficiencies between the GV3101 WT and two *tufAB^Pst4-15^* strains (AS201 and AS202) in *A. thaliana* seedlings by quantitative GUS activity (b) GUS staining (c, scale bar =1 cm). Results are representative of three independent experiments. (d) Comparison of genome editing efficiency between the GV3101 WT and two *tufAB^Pst4-15^* strains (AS201 and AS202) measured by the percentage of tribenuron-resistant T1 plants (left) and number of edited T1 plants per three T0 plants (right). (e) Comparison of stable transformation efficiency by the number of hygromycin-resistant T1 plants per three T0 plants, with no significant differences between WT GV3101 and the two *tufAB^Pst4-15^*strains (AS201 and AS202). Seeds were collected from all flowers. Each data point represents biological replicates from three independent experiments. All values are shown as mean ± SD, with *p*-values calculated by Student’s t-test.

Encouraged by these results, we next tested whether the *tufAB^Pst4-15^*strains could improve genome editing efficiencies in Col-0 by AMT-FI. Compared to GV3101, both AS201 and AS202 significantly enhanced genome editing rates and produced more edited T1 plants (Fig. 6d). As expected, we observed no significant difference in T-DNA integration rates or in the total number of transgenic T1 plants (Fig. 6e).

### Sequencing-based screening of genome-edited plants

Our floral inoculation analysis demonstrated that flowers opened at 6 DPI showed a stable transformation efficiency ∼26% (Fig. 4c). Based on this high efficiency, we reasoned that it may be possible to recover genome-edited T1 plants without prior selection for the T-DNA marker (Fig. 7a).

**Figure 7.**
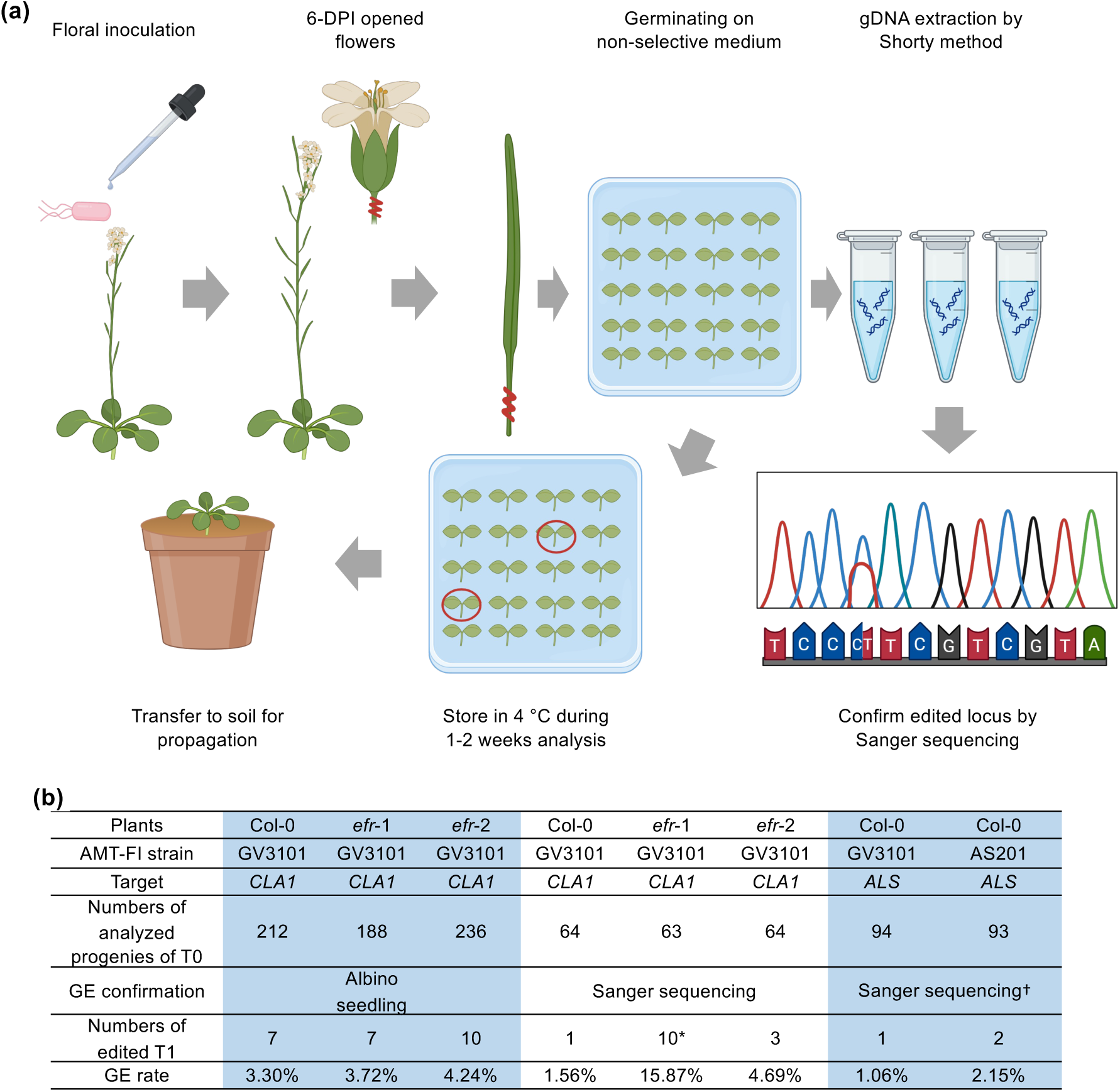
Sequencing-based screening of genome-edited plants. (a) Workflow for screening genome-edited T1 plants from flowers that opened at 6 DPI. Seeds were germinated in 1/2 MS medium with only supplemented with timentin to suppress *Agrobacterium* growth without hygromycin selection. 10-day-old T1 plants were analyzed by Sanger sequencing at the *ALS* or *CLA1* loci. The diagram was created with BioRender.com. (b) The number of sequenced T1 plants and editing rates. ^†^indicates the C to T editing at Pro197 locus conferring tribenuron resistance. Statistically significant differences in genome editing (GE) rates between *efr*-1 and WT Col-0 are indicated (\**p* < 0.001, chi-square analysis).

To test this, we targeted *CLA1* gene, which allows easy identification of edited plants based on their albino phenotype, complemented by sequencing analysis. We germinated T1 seeds from Col-0 and *efr* mutants on MS medium without hygromycin selection. Among these seedlings, we observed biallelically edited albino plants at the frequencies of 3.3% in Col-0, 3.7% in *efr*-1, and 4.2% in *efr*-2. We also identified green seedlings carrying monoallelic edits, identified by Sanger sequencing. Their editing rates were 1.6% in Col-0, 15.9% in *efr*-1, and 4.7% in *efr*-2 (Fig. 7b). To further validate this strategy, we targeted the *ALS* gene without applying tribenuron selection. Among the T1 seedlings, we identified one edited plant out of 94 T1 seedlings using GV3101 and two out of 93 using the *tufAB^Pst4-15^* mutant, AS201. All three edited T1 plants carried C-to-T base editing at the Pro197 locus, consistent with the expected *ALS* editing outcome (Fig. 7b).

In conclusion, genome-edited T1 plants can be obtained without prior selection for T-DNA integrated plants when screening seeds from flowers opened at 6 DPI. Furthermore, evading EFR-mediated PTI enhances genome editing efficiency, further supporting the role of plant immunity in constraining *Agrobacterium*-based genome engineering.

## DISCUSSION

In this study, we provide mechanistic and practical insights into the *Agrobacterium-* mediated floral transformation process. By tracking transformation events with a dual-reporter system, we identified the optimal transformation window, leading to the success in obtaining edited plants without prior selection of T-DNA integration. Our findings also reveal EFR-mediated immunity as an important barrier to efficient transient transformation and genome editing via AMT-FI.

By integrating floral stage selection, immune evasion, and *Agrobacterium* engineering, our study provides a practical guideline to advance *Agrobacterium-* mediated plant transformation and genome engineering. The recommended AMT-FI procedure is summarized in Fig. S6. First, the engineered *tufAB*-modified strain AS201, which showed relatively better performance than AS202, is recommended to replace GV3101 for plant transformation in *Arabidopsis* or other plants possessing *EFR*. Second, we recommend to collect seeds only from flowers that opened 5∼9 day-post-inoculation (DPI), which yield 4∼5% hygromycin-resistant transformants (Fig. 4c, 5), compared with ∼2% when seeds are collected from all flowers (Fig. 6e). Third, for screening edited T1 plants without prior selection for T-DNA integration, collecting seeds exclusively from flowers opened at 6 DPI can reduce labor and cost as well as increase the frequency of obtaining edited plants (Fig. 7). In our experimental condition, it is critical to follow the experimental procedure for growing only one plant in a small pot (5 cm upper diameter × 5 cm height × 3.5 cm bottom diameter). When growing three plants in one larger pot, the peak of AMT-FI efficiency is generally delayed by approximately one day.

### Temporal and spatial tracking of *Agrobacterium* entry and transformation

Using RUBY as a T-DNA reporter, we detected strong RUBY signals in sepal, transmitting tissues, ovules, and developing seeds (Fig. 2). Notably, no RUBY signals were detected in pollen grains, consistent with earlier findings using GUS reporter (Ye *et al*., 1999; Bechtold *et al*., 2000; Desfeux *et al*., 2000). The variation in transformation efficiency likely reflects cell-type specific susceptibility to *Agrobacterium* infection or differential transcriptional activity of the CaMV 35S promoter (Kiselev *et al*., 2021). Additionally, we observed diverse RUBY expression patterns among hygromycin-resistant transgenic plants. Some plants expressed RUBY throughout their life cycle, while others showed expression only during seedling stages or specific organs. This variability may result from positional effects of T-DNA integration site or differences in promoter activity (Gelvin, 2003; Roca Paixao & Deleris, 2024). The presence of hygromycin-sensitive but RUBY-positive seedlings suggests either truncation of the T-DNA at the left border (Fig. 3a), where the hygromycin-resistance cassette is located, or the weaker activity of the *nos* promoter driving expression of the hygromycin-resistance gene.

Although various floral tissues and cell types can be transformed (Figs. 1, 2), only female gametes serve as heritable targets for producing progenies in *Arabidopsis* (Ye *et al*., 1999; Bechtold *et al*., 2000; Desfeux *et al*., 2000). Therefore, *Agrobacterium* entry into the locule of developing gynoecium is a crucial step in successful transformation (Desfeux *et al*., 2000). Our data show that *Agrobacterium* enters the locule of open gynoecium within hours, colonizes reproductive tissues, and persists around the micropyle of ovules, potentially enabling transformation of egg cells and surrounding tissues. Despite early colonization, transformation events in gynoecium cells only occur in open flowers from 5 DPI onward, coinciding with floral opening and micropyle exposure.

Transient T-DNA expression can be observed as early as 1 DPI and highly abundant at 2 DPI when transforming somatic cells like leaves, seedlings, and root explants (Gelvin, 2003; Wu *et al*., 2014; Lopez-Agudelo *et al*., 2024). The delayed expression in germ cells may reflect differences in promoter activity, cell-type susceptibility, or route of *Agrobacterium* entry. Given that pollen tubes access ovules via the micropyle only after floral opening, they may serve as important carriers, facilitating *Agrobacterium* entry into ovules for transformation. Once inside, *Agrobacterium* could associate with male gametes or directly attach to egg cells for T-DNA transfer during fertilization. This proposed mechanism explains the high AMT-FI efficiency observed in flowers opening at 5-9 DPI and suggests that transformation occurs in young floral buds 5-9 days before anthesis, supporting previous findings (Clough & Bent, 1998; Ye *et al*., 1999; Desfeux *et al*., 2000).

### AMT peaks in flowers opened at 6 DPI and negatively correlate with seed viability

Tracking transformation events by fluorescent marker revealed *Agrobacterium* targets integument cells (Fig. 1l, 1m), consistent with previous findings (Bechtold *et al*., 2003). However, only the transformed egg cells contribute to transgene inheritance. This explains why RUBY-based transformation rates are significantly higher than those determined by hygromycin resistance (Figs. 3c, 4c). The use of the RUBY reporter enabled the discovery of floral stages accessible for AMT-FI, revealing that nearly all ovaries from flowers open at 6 DPI were transformed (Fig. 3b), yielding averaged an 26% stable Hyg^R^ T1 progeny (Fig. 4c). However, high transformation rates were accompanied by increased ovule abortion, resulting in fewer viable seeds (Figs. 3d, 4a, 4c). In contrast, *efr* mutants exhibited lower abortion rates and higher seed viability per silique, suggesting that EFR-triggered immunity contributes to transformation-induced ovule loss. Evading plant immune responses could therefore enhance the efficiency of transgenic seed production via AMT-FI.

### Evasion of EFR-mediated immunity enhances genome editing efficiency but not stable transformation

Although EFR-mediated immunity was discovered nearly two decades ago (Zipfel *et al*., 2006), its influence on stable transformation and genome editing remain unclear until recent studies. While *efr*-1 mutants showed no improvement on gene targeting or stable transformation efficiencies (van Tol *et al*., 2022), they do exhibit enhanced transient expression (Zipfel et al., 2006; Wu, Liu et al., 2014; Yang et al., 2023). Our results revealed higher RUBY transformation rates in *efr* ovules, and confirmed no significant increase in stable transformation measured by hygromycin resistance (Figs. 4c, 6e).

However, we showed that genome editing efficiencies were higher in *efr* mutants and in Col-0 WT inoculated with the two engineered GV3101::*tufAB^Pst4-15^*strain, AS201 and AS202 (Figs. 5, 6). This suggests that genome editing can be decoupled from T-DNA integration. This is further supported by evidence that genome-edited plants were successfully obtained from the *teb-5* (*polQ)* mutant, whereas T-DNA integrated plants were not (van Tol *et al*., 2022). Thus, we proposed that enhancement of editing efficiencies may result from higher transient expression of CRISPR/Cas9 in egg cells in the absence of EFR-triggered immunity. Moreover, because editing can occur without DNA integration (e.g., via ribonucleoprotein delivery), it may serve as a proxy for measuring transgene expression levels in germline cells.

### Innovative strategies for enhancing genome editing efficiency

Precision editing is critical for inducing dominant phenotypes by targeting an endogenous gene. However, current methods exhibit low efficiency compared to mutagenesis by Cas9 nuclease (Huang & Puchta, 2021). To overcome these limitations, we used an improved A3A-CBE base editor and achieved a high editing-to-integration ratio (∼0.2) (Fig. 5). This construct, together with our optimized screening workflow, enables efficient dissection factors that influence stable transformation and genome editing efficiency in *Arabidopsis*.

Several recent studies have introduced innovative approaches to enhance stable transformation and genome editing. These include engineering of *Agrobacterium* strains to deliver type III secretion system effectors, T-DNA concatenation, and vector copy number manipulation (Raman *et al*., 2022; Dickinson *et al*., 2023; Szarzanowicz *et al*., 2024). Our success in engineering *Agrobacterium* strains with modified EF-Tu to evade EFR recognition, significantly enhanced genome editing efficiency. This reinforces the idea that boosting transient gene expression is a key determinant for successful genome modification via CRISPR/Cas9.

Despite native EFR is specific to Brassicaceae, several non-Brassicaceae crops have been engineered to express EFR for disease resistance (Lu *et al*., 2015; Schwessinger *et al*., 2015; Boschi *et al*., 2017; Kunwar *et al*., 2018; Piazza *et al*., 2021). These transgenic plants may show reduced *Agrobacterium-*mediated transformation. Our engineered GV3101::*tufAB^Pst4-15^* strains could overcome this limitation. Beyond EF-Tu, other *Agrobacterium-*derived PAMPs such as translation initiation factor IF1, cold-shock protein, and flagellin also trigger PTI via distinct receptors in different species (Saur *et al*., 2016; Fürst *et al*., 2020; Fan *et al*., 2022). This work sets the stage for the development of multi-modified *Agrobacterium* strains with broad compatibility, offering a roadmap to enhance AMT across diverse plant taxa.

## AUTHOR CONTRIBUTIONS

M.-S.L, T.-K.H., C.-H. K. and E.-M.L, contributed to the conceptualization of the project. M.-S.L. designed and performed AMT-FI. T.-K.H. designed and performed the genome editing assay. M.-S.L., T.-K.H., Y.-C.W. and S.-C.W. performed experiments and analyzed data. M.-S.L., T.-K.H., and E.-M.L. wrote the initial manuscript. E.-M.L., C.-H. K. and C.-H.W. supervised the project. M.-S.L and T.-K.H contributed equally. All authors reviewed and edited the manuscript.

## ACKNOWLEDGEMENTS

The authors acknowledge the members of the Lai, Kuo, and Wu labs for discussion and suggestion. We also thank the technical assistance of Live Cell Imaging Core Lab and Sanger DNA sequencing service provided by Genomic Technology Core located at the Institute of Plant and Microbial Biology, Academia Sinica. The funding was supported by the Academia Sinica Grand Challenge Program (AS-GC-111-L02) to E.-M.L., C.-H. K. and C.-H.W, and the intramural fund of the Institute of Plant and Microbial Biology, Academia Sinica.

## SUPPORTING INFORMATION

Table S1. Plasmid information

Table S2. Primer list

Figures S1-S6

## DATA AVAILABILITY

The data supporting the findings of this study are available within the article or as Supporting Information. Strains and plasmids generated in this study are available either on Addgene (no. 234350-234353, 237864-237866) or upon request (Table S1)

---

**Figure S1.**
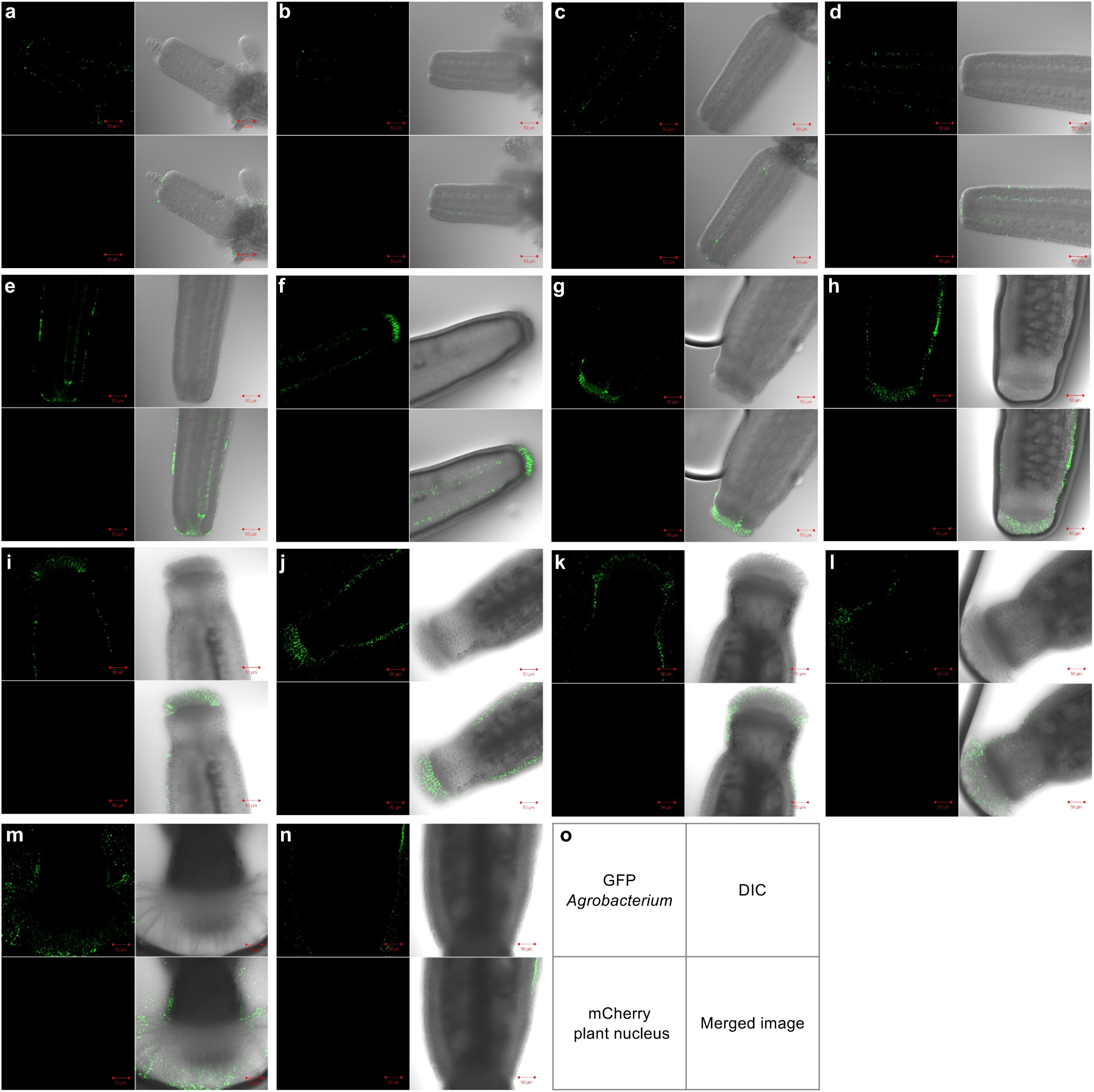
Basipetal analysis of *Agrobacterium* distribution along *Arabidopsis* inflorescence at 1 DPI. A basipetal series of flowers from the inflorescence tip (a) to the first open flower at the base (m and n) were dissected to visualize distribution of *Agrobacterium* expressing GFP reporter at 1 DPI. The order was determined in considering the position and developmental stage of flowers. Each figure consists of four imaging channels (o). Scale bars indicated 50 µm. Panels a-e show open gynoecia, f-n are closed gynoecia, m and n are from different regions of the same flower.

**Figure S2.**
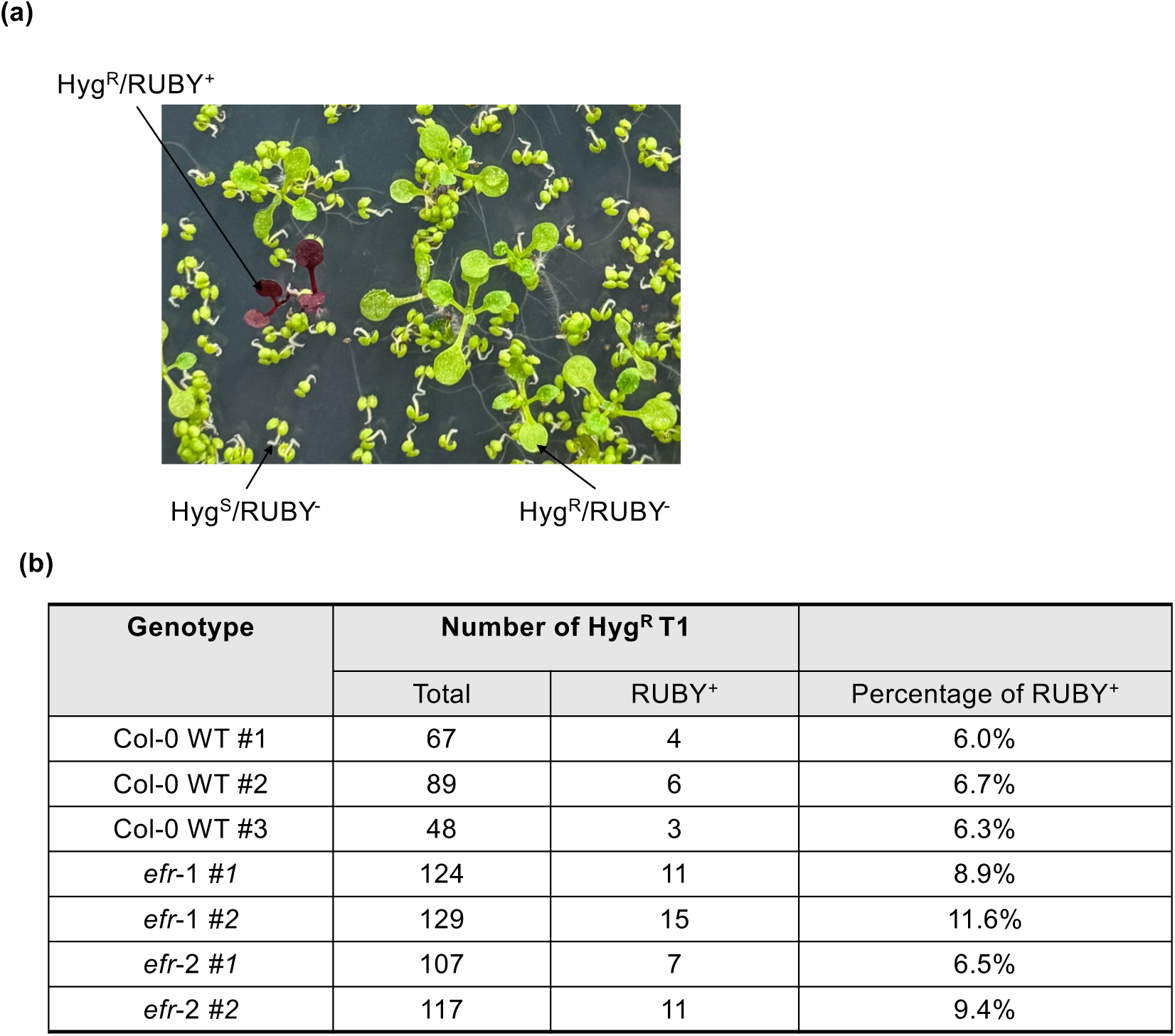
Number and percentage of Hyg^R^/RUBY^+^ seedlings. (a) Representative photo of T1 seedlings screening on hygromycin containing 1/2 MS plate. The hygromycin-resistant T1 exhibiting RUBY signals of whole seedling (Hyg^R^/RUBY^+^) was indicated, among the hygromycin-resistant green and larger T1 seedling (Hyg^R^/RUBY^-^), and hygromycin-sensitive green but small seedlings (Hyg^S^/RUBY^-^). The hygromycin-sensitive T1 seedlings with RUBY signals (Hyg^S^/RUBY^+^) shown in Fig. 2j are rarely occurred and not present in this photo. (b) Number and percentage of Hyg^R^/RUBY^+^ T1 seedlings in Col-0 WT and *efr* mutants. Total number of Hyg^R^ T1 seedlings (Total) and those exhibiting RUBY signals of whole seedling (RUBY^+^) was indicated and the percentage of Hyg^R^/RUBY^+^ T1s among total Hyg^R^ T1s.

**Figure S3.**
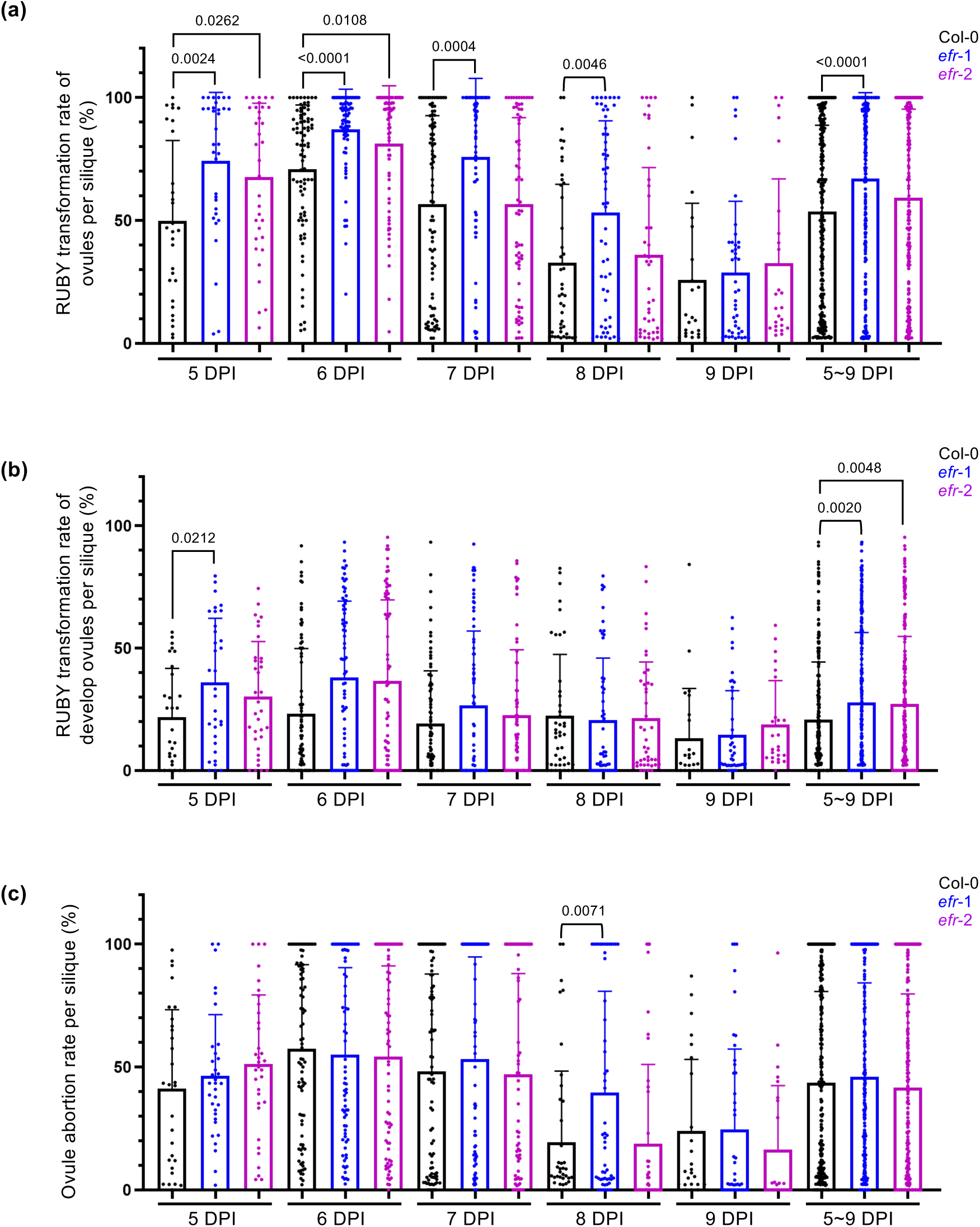
EFR-mediated immunity exerts negative impacts on ovule development and RUBY-based transformation rate. GV3101 carrying the 35S::RUBY construct was used for AMT-FI to assess RUBY-based transformation efficiency and ovule abortion rate in Col-0 WT and *efr* mutants. Siliques from flowers that opened at 5-9 DPI were analyzed to count the total number of ovules, the number of developed ovules with or without RUBY signals, and the number of abortive ovules with or without RUBY signals per silique. (a) Daily RUBY-based transformation rate per silique. The percentage of RUBY-positive ovules (including both developed and abortive ovules) per silique was calculate at each DPI, as well as cumulatively across 5-9 DPI. Each data point represents the percentage of RUBY-positive ovules per silique. (b) Daily RUBY-based transformation rate of developed ovules per silique. The percentage of RUBY-positive developed (non-abortive) ovules per silique was measured at each DPI, as well as cumulatively across 5-9 DPIs. Each data point represents the percentage of RUBY-positive developed ovules per silique. (c) Daily ovule abortion rate per silique. The percentage of abortive ovules (including both RUBY-positive and RUBY-negative) per silique was recorded at each DPI and cumulatively across 5-9 DPI. Each data point represents the percentage of abortive ovules per silique. Data are shown in mean ± SD from three independent experiments, with *p*-values calculated by Student’s t-test between Col-0 WT and *efr* mutants. Related to Figure 3.

**Figure S4.**
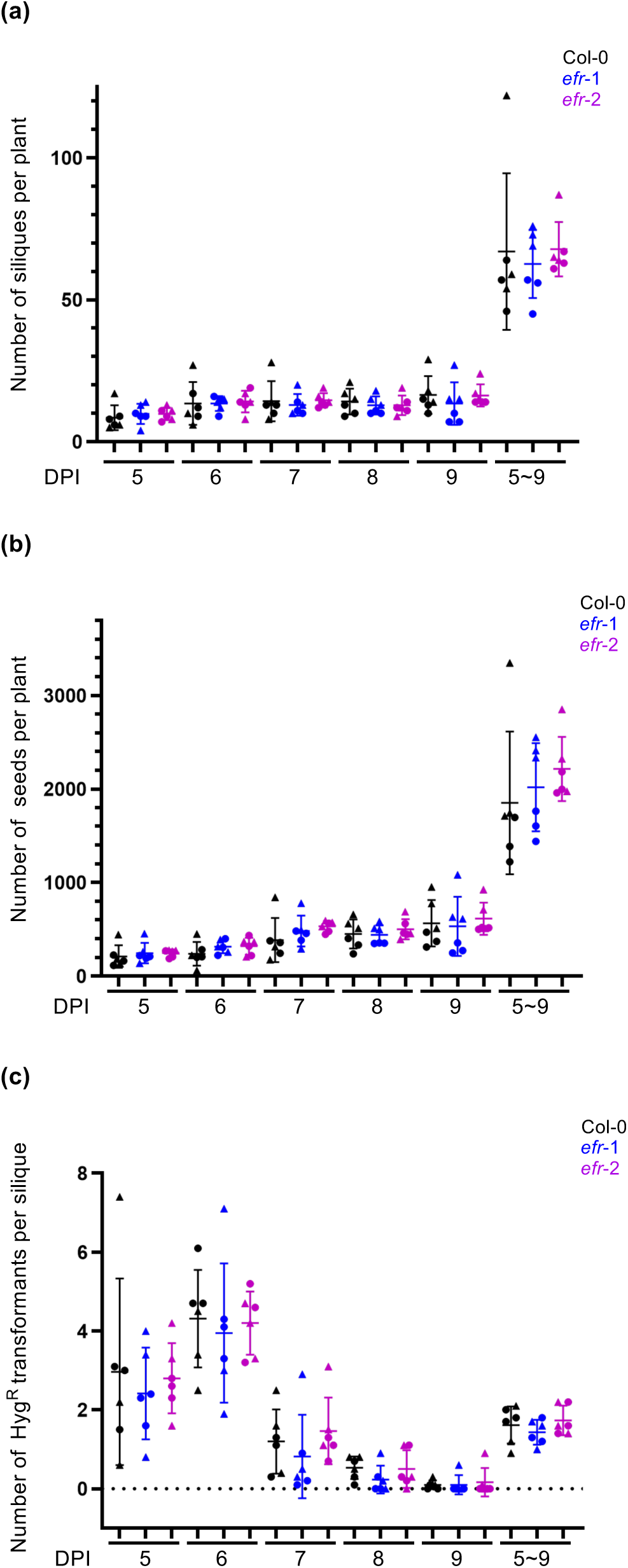
Number of siliques, seeds, and hygromycin-resistant seedlings in Col-0 WT, *efr*-1 and *efr*-2 mutants after AMT-FI. Siliques and seeds developed from flowers that opened at 5-9 DPI in Col-0 WT, *efr*-1 and *efr*-2 mutants were analyzed through number of siliques (a) and seeds (b) per plant, and averaged number of hygromycin-resistant (Hyg^R^) seedlings per silique (c). Data are mean ± SD of six biological replicates from two independent experiments (circle and triangle symbols).

**Figure S5.**
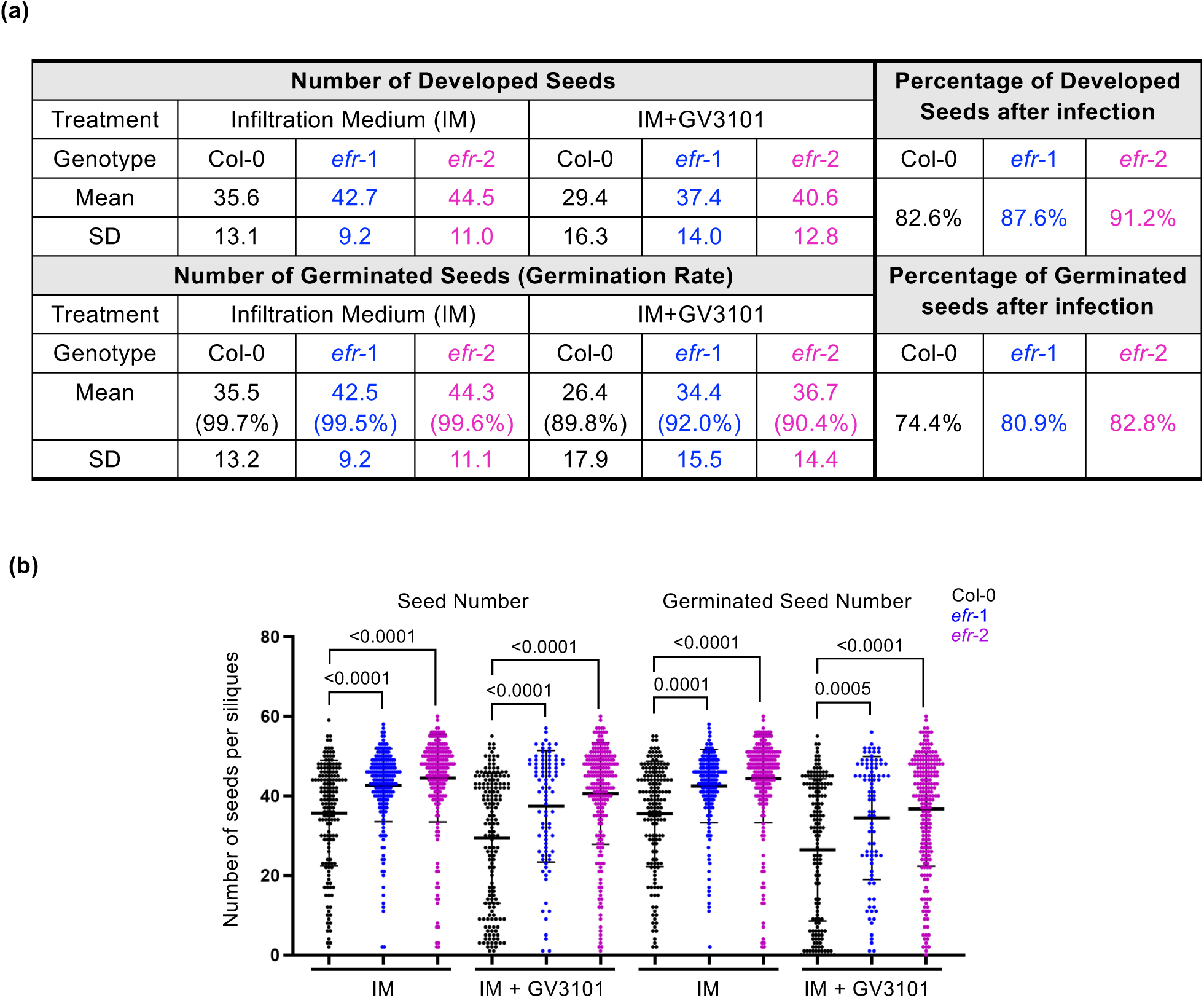
*Agrobacterium* infection and EFR-mediated immunity affect seed outputs. (a) Table showing seed number per silique from plants treated with infiltration medium alone or with *Agrobacterium.* Percentage of developed/germinated seeds after infection was normalized to those from infiltration medium (IM) treatment. The resulting percentage of developed and germinated seeds indicated the impact on seed viability. (b) Seed counts and germination rates following IM alone or with *Agrobacterium* IM (IM + GV3101) treatments. Siliques were collected from flowers that opened at 5–9 DPI. The total number of seeds and the number of germinated seeds were compared between treatments. Data were collected from two plants per genotype, except for *efr*-1 infected with GV3101, where data were obtained from one plant. Data are shown as mean ± SD, with *p*-values calculated using Student’s *t*-test to compare Col-0 WT and *efr* mutants.

**Figure S6.**
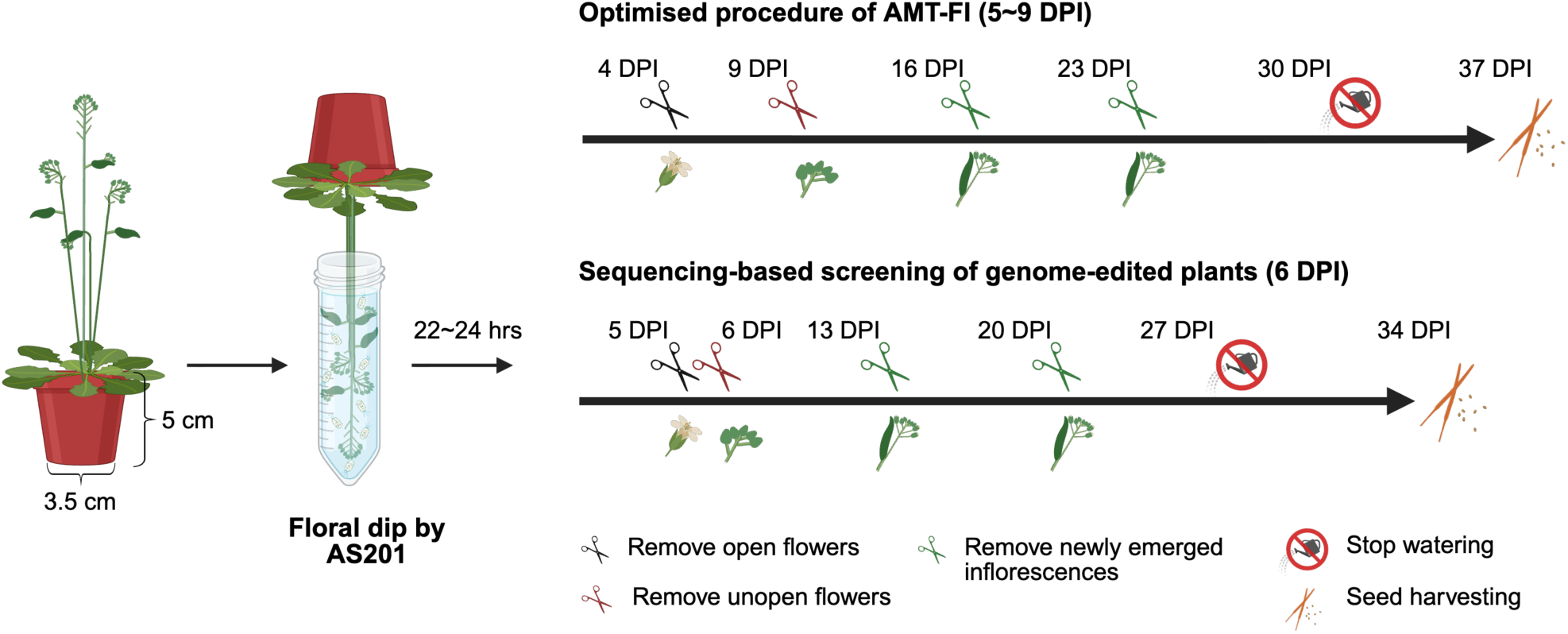
The recommended AMT-FI procedure. The experimental procedures for achieving higher AMT-FI efficiency and for screening genome-edited T1 plants without prior selection for T-DNA integration are illustrated. The engineered *tufAB*-modified strain AS201 is recommended for plant transformation in *A. thaliana* or other plant species possessing *EFR*. The optimized transformation window was determined under the growth conditions of *A. thaliana*, with each plant cultivated in a small pot (5 cm upper diameter × 5 cm height × 3.5 cm bottom diameter). The diagram was created with BioRender.com.

**Supplemental Table 1.** Strain and Plasmid information.

| Lab collection # | ***Agrobacterium* strain/** Plasmid/<br>Antibiotics resistance | Characteristics | Source |
| --- | --- | --- | --- |
| EML485 | **GV3101**/Rif<sup>R</sup>Gen<sup>R</sup> | GV3101 containing pMP90, which is pTiC58 derivative with T-DNA deletion and gentamicin resistance gene | Koncz and Schell, 1986 |
| EML8561 | **AS201**/ Rif<sup>R</sup>Gen<sup>R</sup> | GV3101::*tufAB*<sup>Pst4-15</sup> mutant by modifying *tufB* followed by *tufA*, with both genes encoding a chimeric EF-Tu with amino acids 4-15 from corresponding *Pst* EF-Tu | This study |
| EML8559 | **AS202**/ Rif<sup>R</sup>Gen<sup>R</sup> | GV3101::*tufAB*<sup>Pst4-15</sup> mutant by modifying *tufA* followed by *tufB*, with both genes encoding a chimeric EF-Tu with amino acids 4-15 from corresponding *Pst* EF-Tu | This study |
| EML3375 | pRL662-GFP(S65T)/ Gen<sup>R</sup> | Broad host range plasmid confers gentamycin resistance and expression of GFP(S65T) | Yu et al., 2021 |
| EML6428 | pRL662-GFP(S65T)/Spe<sup>R</sup> | Broad host range plasmid confers spectinomycin resistance and expression of GFP(S65T) | This study, Addgene#234350 |
| EML6409 | 35S::RUBY/Spe<sup>R</sup> | Binary vector carrying T-DNA expressing RUBY reporter and hygromycin resistance | Addgene#160908 |
| EML6442 | 35S:mCherry2-NLS/Kan<sup>R</sup> | Binary vector carrying T-DNA expressing mCherry2 florescent proteins | This study, Addgene#234351 |
| EML8357 | pA3A-CBECas9/Kan<sup>R</sup> | Binary vector carrying T-DNA expressing base editor driven by egg-cell specific promoter | This study, Addgene#234352 |
| EML8359 | pA3A-CBECas9-Pro197/Kan<sup>R</sup> | Binary vector carrying T-DNA expressing base editor targeting *ALS Pro197* locus | This study, Addgene#234353 |
| EML8532 | pA3A-CBECas9-CLA1/Kan<sup>R</sup> | Binary vector carrying T-DNA expressing base editor targeting *CLA1*, generating early stop codon | This study |
| EML8533 | pA3A-CBECas9-PDS3/Kan<sup>R</sup> | Binary vector carrying T-DNA expressing base editor targeting *PDS3*, generating early stop codon | This study |
| EML121 | pJQ200KS/Gen<sup>R</sup> | Suicide plasmid contains *sacB* and Gm<sup>R</sup> *traJ oriT* for the selection of double crossover events | Quandt and Hynes, 1993 |
| EML8543 | pEML041003/Kan<sup>R</sup> | pJQ200KS derived suicide vector for gene replacement in *Agrobacterium* | This study, Addgene#237864 |
| EML8552 | pEML041003-tufAPst4-15/ Kan<sup>R</sup> | Suicide vector for replacing 4<sup>th</sup> to 15<sup>th</sup> amino acids of *tufA* | This study, Addgene#237865 |
| EML8553 | pEML041003-tufBPst4-15/ Kan<sup>R</sup> | Suicide vector for replacing 4<sup>th</sup> to 15<sup>th</sup> amino acids of *tufB* | This study, Addgene#237866 |

Gen<sup>R</sup>, gentamycin resistance; Kan<sup>R</sup>, kanamycin resistance; Rif<sup>R</sup>, rifampicin resistance; Spe<sup>R</sup>, spectinomycin resistance

**Supplemental Table 2.**
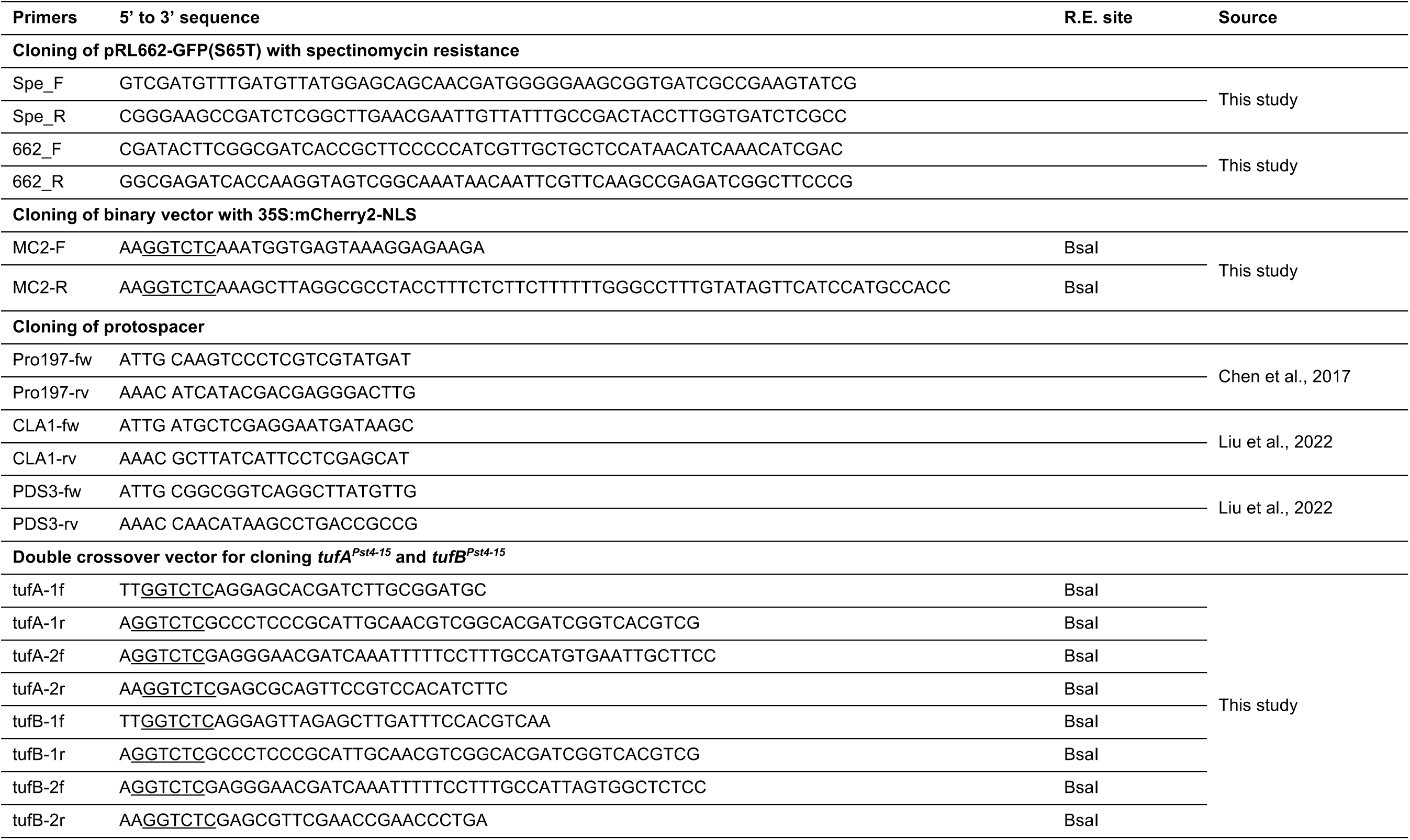
Primer information.

